# Post-translational toxin–antitoxin control and RecBCD surveillance underlie phage defense in a constitutively armed Type VI retron

**DOI:** 10.64898/2026.04.27.721009

**Authors:** Fernando M. García-Rodríguez, Francisco Martínez-Abarca, Rafael Molina, Nicolás Toro

## Abstract

Type VI retrons are bacterial defense systems that couple reverse transcription to phage immunity through dedicated toxin–antitoxin modules, yet the mechanisms linking ssDNA synthesis to toxin activation remain unclear. Here we characterize Retron-Sen3, a Type VI retron from *Salmonella enterica* encoding a small protein (SP) toxin, an HTH antitoxin, and a reverse transcriptase (RT). Unlike previously examined Type VI retrons, Retron-Sen3 constitutively produces RT-dependent ssDNA species in the absence of phage infection, and these levels remain unchanged during infection, establishing that defense activation is uncoupled from ssDNA synthesis. Comparative analyses revealed a conserved ncRNA architecture across the cl2075 lineage, including a previously undescribed stem–loop essential for defense, and an extended transcript encompassing the SP coding region within a structurally conserved RNA framework. SP is constitutively produced and intrinsically toxic, whereas HTH neutralizes SP toxicity, is required for basal viability, and physically associates with RT *in vivo* independently of catalytic activity; notably, the HTH–SP toxin–antitoxin module operates independently of RT and ncRNA. Phage escape mutants and genetic analyses establish that inhibition of the host RecBCD complex triggers defense activation, identifying RecBCD surveillance as the signal that couples phage infection to toxin activation. Together, these findings reveal a post-translational regulatory mechanism distinct from the msDNA-dependent translational control described for Retron-Vpa2, establishing mechanistic diversity within this retron family.

## Introduction

Bacteria encode a remarkable diversity of defense systems that protect against bacteriophage infection (Bernheim and Sorek, 2020; Doron et al., 2018; Gao et al., 2020). Among these, retrons have recently emerged as widespread tripartite defense systems composed of a reverse transcriptase (RT), a structured non-coding RNA (ncRNA), and an associated effector protein with diverse predicted activities (Mestre et al., 2020; Millman et al., 2020; Bobonis et al., 2022). Initially discovered as enigmatic retroelements, retrons are now recognized as anti-phage systems that typically trigger abortive infection, wherein the infected cell sacrifices itself to limit viral spread through the population (Millman et al., 2020; Lopatina et al., 2020). In canonical retrons, the ncRNA contains contiguous msr and msd regions that form the template for reverse transcription, generating multicopy single-stranded DNA (msDNA), an unusual branched DNA–RNA molecule (Inouye et al., 1989; Lampson et al., 1989; Hsu et al., 1992) that plays central regulatory roles in defense. Beyond their defensive role, retrons have been harnessed as powerful genome-editing tools in both prokaryotes and eukaryotes (Simon et al., 2019; Farzadfard and Lu, 2014; Bhattarai-Kline et al., 2022; Lopez et al., 2022; Fishman et al., 2025).

Retrons are currently classified into 13 types based on the phylogeny of their encoded RTs and the predicted structural and functional features of their associated effectors (Mestre et al., 2020). In many retrons, the msDNA produced by the RT acts as a molecular sentinel of the cellular state: its structural integrity signals the absence of phage infection, whereas its modification or depletion relieves effector inhibition and triggers defense (Millman et al., 2020; Bobonis et al., 2022; Gao et al., 2020). Retron effectors are highly diverse and include nucleases, deaminases, NAD⁺-hydrolases, and other enzymatic activities that, once activated, arrest bacterial growth or cause cell death (Gao et al., 2020; Carabias et al., 2024). A recurring effector strategy involves toxin–antitoxin (TA) modules, in which the RT–msDNA complex acts as an antitoxin that directly neutralizes a toxic effector. Phage-encoded triggers disrupt this complex, releasing or activating the toxin to abort phage replication (Bobonis et al., 2022; Wang et al., 2022; García-Rodríguez et al., 2025). This logic parallels the broader bacterial immune strategy in which constitutively expressed antitoxins neutralize their cognate toxins until infection-induced signals activate them to arrest cell growth (Lopatina et al., 2020; LeRoux and Laub, 2022).

For several retron types, including Types VI, VII-A1, VII-A2, VIII, X, XI, and XIII, the structured msr–msd sequences of the ncRNA have not yet been identified (Mestre et al., 2020). The Type VI retron system encodes RTs belonging to clade 3 and has a characteristic genomic architecture, typically encoding a Cro/C1-type HTH protein and, in a subset, a small accessory SP protein from one of four conserved clusters (cl2075, cl2760, cl3272, and cl3897), defining the canonical SP–HTH–RT organizational arrangement (Mestre et al., 2020). In a systematic survey of all 13 retron classes, Type VI retrons failed to produce detectable ssDNA species under standard expression conditions (Khan et al., 2025), a result later confirmed for two additional Type VI members expressed as complete operons under native promoters (Zhang et al., 2025). Recent characterization of Retron-Vpa2, a Type VI retron from *Vibrio parahaemolyticus*, revealed a fundamentally different logic from the canonical retron TA model: rather than acting as an antitoxin that neutralizes a toxic effector, the msDNA acts as a positive activator of toxin translation upon phage infection, suggesting that Type VI retrons may employ mechanistic principles distinct from those of other retron types (Zhang et al., 2025).

Here we characterize Retron-Sen3, a Type VI retron from *Salmonella enterica* belonging to the well-represented cl2075 lineage within Proteobacteria. We show that Retron-Sen3 constitutively produces RT-dependent ssDNA species in the absence of phage infection, distinguishing it from all previously characterized Type VI retrons, and that defense activation occurs without detectable changes in ssDNA abundance. Functional analyses further reveal that the encoded SP toxin is constitutively produced and restrained by a cognate HTH antitoxin, establishing a post-translational toxin-antitoxin regulatory module. Phage escape mutants and host genetics establish that inhibition of the RecBCD pathway is the signal that triggers defense activation. Together, these findings reveal a previously unrecognized regulatory strategy within the Type VI retron family and provide a mechanistic framework for understanding how retron-mediated defense can operate independently of msDNA-dependent translational control.

## Material and Methods

### Bacteria, phage strains and growth conditions

Bacteriophages T3, T5, T7, λ, M13, Qβ, and φX174, and *E. coli* strains LE392, Hfr 3000 U432, MG1655, B, and BW25113 were obtained from the DSMZ–German Collection of Microorganisms and Cell Cultures GmbH (Braunschweig, Germany). Phages T2, T4, P1, MS2, and φV-1 were obtained from the American Type Culture Collection (ATCC, Manassas, VA, USA). The completed BASEL collection (Maffei et al., 2021; Humolli et al., 2025), comprising 107 phages isolated from environmental samples and infecting *E. coli* as a model organism, was kindly provided by Dr. Alexander Harms.

*E. coli* MG1655 was used as the host strain for phages T2, T4, T5, T7, P1, and φV-1. Phages M13, MS2, Qβ, and φX174 were propagated on strain Hfr 3000 U432. Assays with phages from the completed BASEL collection were performed using *E. coli* K-12 MG1655 ΔRM and *E. coli* K-12 BW25113 Δ*gtrS wbbL(+)*. Phage λ was assayed on strain LE392, and phage T3 on *E. coli* B. The *E. coli* BW25113 single-gene deletion mutants Δ*recB*, Δ*recC*, and Δ*recD* were obtained from the Keio collection (Baba et al. 2006)

When required, ampicillin and chloramphenicol were used at final concentrations of 100 µg/ml and 200 µg/ml, respectively.

### Plasmid construction

Plasmid used in this study are listed in **Table S1**. The Retron-Sen3 locus from *S. enterica* was synthesized by GenScript Corp. (Piscataway, NJ, USA) and provided cloned into pUC57mini, generating the corresponding ampicillin-resistant plasmid p57m VI. A derivative of p57m VI carrying the DD193–194AA substitution in the RT catalytic site (p57m VI RTmut), its C-terminally 3×FLAG-tagged version (p57m VI RTmut-F), the wild-type RT C-terminally tagged with 3×FLAG (p57m VI RT-F), and the SP protein C-terminally tagged with 3×FLAG (p57m VI SP-CF) were also synthesized by GenScript from p57m VI. Frameshift mutants of SP and HTH were generated by inserting a guanine nucleotide after position +36 of their respective coding sequences, yielding both untagged variants (p57m VI SPmut and p57m VI HTHmut) and C-terminally 3×FLAG-tagged versions (p57m VI SPmut-CF and p57m VI HTHmut-CF); all four constructs were synthesized by GenScript. Two additional derivatives targeting the 5′-conserved stem–loop of the ncRNA were also generated by GenScript: p57m VI Δ18nt-SL, carrying a deletion of the 18 nucleotides comprising the 5′-conserved stem–loop, and p57m VI 18nt-SLmut, in which positions 1, 3, 5, and 8 of the 5′-conserved stem–loop were substituted with adenine (position 1,3 and 5) or thymine (position 8). The plasmid p57m VI ΔncRNA was generated by deleting the SL3 region of the ncRNA from p57m VI.

For the expression of SP-HTH, and their respective mutant variants, the corresponding coding sequences were cloned into the arabinose-inducible expression plasmid pBAD33 between the *Kpn*I and *Sal*I restriction sites using the NEBHiFi Assembly kit (New England Biolabs) according to the manufacturer’s instructions. The inserts were amplified by PCR using primers HF-SPHTH-Rv (CAAGCTTGCATGCCTGCAGGTCGACCTATATGAGGGCGC) and HF-SPHTH-Fw (AGCGAATTCGAGCTCGGTACCATGAAGTACGAAACTTTG), with plasmids p57m VI, p57m VI-SPmut, and p57m VI-HTHmut serving as templates for the wild-type and the SPmut-HTH and SP-HTHmut mutant constructs, respectively. The individual genes of the phage λ Red operon (*gam*, *beta*, and *exo*), as well as the *beta*+*exo* and *gam*+*beta*+*exo* combinations, were placed under the control of the arabinose-inducible promoter in pBAD33 and synthesized by GenScript Corp.

### RNA-seq analysis

Total RNA was extracted from E. coli cells expressing the complete Retron-Sen3 module, depleted of ribosomal RNA, and subjected to non-strand-specific Illumina sequencing. Sequencing reads were mapped to the plasmid sequence containing the Retron-Sen3 locus using Geneious (Biomatters Ltd., Auckland, New Zealand). For plasmid-wide analysis, mapped reads were further processed using custom Python scripts. Coverage at each nucleotide position was calculated from aligned reads and normalized as reads per million (RPM) relative to the total number of reads mapping to the plasmid in each sample. Coverage profiles were smoothed using a rolling average with a 200-nt window to facilitate visualization. The boundaries of the enriched region were defined as positions where the smoothed coverage exceeded 20% of the maximum signal within the retron locus, a criterion applied to avoid inclusion of secondary coverage peaks arising from RPM normalization in distal plasmid regions. Boundaries are indicated by dashed vertical lines in **Supplementary Figure S1**.

### Conservation scoring and sliding-window analysis

Nucleotide sequences corresponding to cl2075-associated Type VI retrons were aligned using MAFFT (Katoh and Standley, 2013) with the --globalpair and --maxiterate 1000 parameters to optimize global alignment accuracy. Per-site conservation scores were computed from the resulting multiple sequence alignment using custom Python scripts (Biopython (AlignIO) and NumPy). For each alignment column, the gap fraction was calculated as the proportion of gap characters relative to the total number of sequences in the alignment. Among non-gap residues, the identity fraction was defined as the frequency of the most abundant nucleotide divided by the total number of non-gap residues in that column. The final conservation score for each position was calculated as:

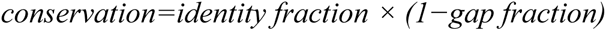

This metric penalizes both sequence variability and gap frequency, yielding conservation values bounded between 0 (fully variable or gap-dominated) and 1 (fully conserved without gaps). To visualize regional conservation patterns, per-site scores were smoothed using a 100-nt sliding window implemented as a moving average (convolution with a uniform kernel). The sliding-window profile was plotted using Retron-Sen3 as the reference sequence. Mean and standard deviation of conservation were calculated for predefined genomic intervals based on alignment positions.

### Structure-guided alignment and structural scanning

To identify conserved structured RNA elements upstream of the SP gene in cl2075-associated Type VI retrons, we initially analyzed ∼450 nt upstream of the SP start codon using structure-guided multiple sequence alignment with mLocARNA (Will et al., 2007). A sliding-window structural scan was performed using 250-nt windows with a 2-nt step size. For each window, truncated sequences from all retrons were aligned using mLocARNA with RNAplfold-based local pairing probabilities (--plfold-span 80, --min-prob 0.05). Consensus secondary structures (SS_cons) were extracted from Stockholm alignments, and a stability ratio was calculated as the fraction of paired positions within each window. Windows exhibiting base-pairing near both termini (≥3 paired positions within the first and last 10 nt) were considered candidate structured elements, consistent with the presence of conserved inverted repeats. Subsequent structural and comparative analyses were restricted to the interval within ∼280 nt upstream of the SP start codon identified by this scan.

### Secondary structure prediction and visualization

The minimum free-energy (MFE) secondary structure and base-pairing probabilities of the Retron-Sen3 ncRNA and full-length extended transcript were predicted using RNAfold from the ViennaRNA package (Lorenz et al., 2011), with partition function calculations performed using the -p option. Base-pairing probabilities were mapped onto the predicted centroid structure for visualization. Graphical representations were generated using forna (Kerpedjiev et al., 2015). Functional elements including stem–loops (SL1–SL6), inverted repeats (IR1, IR2), ribosome-binding sites, and coding sequence boundaries were annotated using custom color assignments within forna.

### Covariance model construction and consensus secondary structure inference

Curated ncRNA sequences corresponding to cl2075-associated Type VI retrons were aligned using mLocARNA (Will et al., 2007) with structure-informed multiple sequence alignment and formatted in Stockholm format. The final alignment comprised 33 representative sequences spanning the conserved ncRNA locus. A covariance model (CM) was constructed from this alignment using cmbuild from the Infernal package (Nawrocki and Eddy, 2013), which encodes both primary sequence conservation and secondary structure constraints. The resulting CM was calibrated using cmcalibrate to enable accurate statistical scoring and E-value estimation during homology searches. The calibrated CM (.cm file) was subsequently used to scan the genomes of cl2075-associated Type VI retrons using cmsearch, thereby confirming the presence and boundaries of homologous structured RNA elements across the dataset. Consensus secondary structure was inferred using the CaCoFold algorithm (Rivas, 2020), which integrates phylogenetic covariation signals while enforcing structural consistency constraints. Covarying base pairs were identified using an E-value threshold of 0.05. In the resulting model, 92 base pairs were predicted, with an average of 5.7 substitutions per base pair. A total of 26 covarying base pairs were observed, compared to an expected 13.5 ± 2.8 under the null model (PPV = 100%; best E = 1.52×10⁻⁸). To assess structural conservation of the extended transcript spanning the IR1–IR2 interval, sequences from 52 cl2075-associated retrons were aligned using mLocARNA (--plfold-span 400, --min-prob 0.03, --indel -6, - -indel-open -18) and subjected to CaCoFold analysis. In the resulting consensus structure, 144 base pairs were predicted, with 18 statistically significant covarying base pairs observed compared to an expected 10.6 under the null model (PPV = 100%; best E = 6.50×10⁻⁸). The IR1–IR2 long-range stem was present in the CaCoFold consensus structure but showed no statistically significant covariation.

### Compensatory substitution analysis

To assess the conservation of the IR1–IR2 long-range stem and the 5′-conserved stem–loop, compensatory substitutions were identified by manual inspection of structure-guided alignments of 53 cl2075-associated retrons. For each stem position, the nucleotide pairs present across all sequences were classified as Watson-Crick, G:U wobble, or mismatch. Positions where two or more distinct Watson-Crick base pairs were observed were classified as compensatory, indicating maintenance of base-pairing potential despite sequence variation. For the IR1–IR2 stem, positions 5, 11 and 14 (numbered from the 5′ end of IR1) were identified as compensatory, each showing Watson-Crick pairs in ≥75% of sequences with multiple pair types. For the 5′-conserved stem–loop, compensatory substitutions were identified at positions 1, 3, 4 and 5 of the 5-bp stem. Alignments were extracted from the MAFFT multiple sequence alignment of the extended transcript interval using custom Python scripts.

### Phage plaque assays

Double-layer plaque assays were performed as described below. *E. coli* host strains were grown to stationary phase in LB at 37 °C. Aliquots of 200 µl were mixed with 10 ml of top agar (LB containing 0.5% agar, 5 mM MgSO₄, and the appropriate antibiotic) and poured onto 12 × 12 cm LB agar plates supplemented with 5 mM MgSO₄ and the corresponding antibiotic. Ten-fold serial dilutions of phage stocks were prepared in LB, and 10 µl of each dilution was spotted onto the plates. Plaques were enumerated after overnight incubation at 37 °C. When individual plaques were too small to quantify, the most concentrated dilution at which no visible plaques were observed was recorded as containing only one plaque.

### Isolation and amplification of mutant phages

To isolate phages capable of escaping Retron-Sen3 defense, phage λ was plated on a lawn of *E. coli* expressing the Retron-Sen3 system using a double-layered plaque assay. Individual plaques (n = 5) were picked into 100 µL of LB and incubated at room temperature for 1 h with occasional vortexing to release phage particles from the agar. Samples were centrifuged at 3,200 × *g* for 10 min to remove agar debris and bacterial cells, and the supernatants were transferred to fresh tubes for analysis. The ability of the recovered phages to escape Retron-Sen3 was evaluated using plaque assays with both a Retron-Sen3–expressing strain and an isogenic strain lacking the system as a control. Ten-fold serial dilutions were prepared for the ancestral phage used in the initial assay and for each of the recovered isolates. Phages exhibiting reduced susceptibility to Retron-Sen3 relative to the ancestral phage were further propagated by picking a single plaque formed on Retron-Sen3–expressing cells into 10 ml of an LB culture (supplemented with 5 mM MgSO₄) of Retron-Sen3–harbouring cells grown to an OD₆₀₀ of 0.3. The cultures were incubated for 6 h at 37 °C with shaking (175 rpm). The resulting lysates were clarified by centrifugation at 3,200 × *g* for 10 min, and the supernatants were filtered through 0.2-µm filters.

### Phage sequencing and analysis of mutants

Phage DNA was extracted from 500 µl of high-titer λ lysate (>10⁷ PFU ml⁻¹). Lysates were treated with DNase I (20 µg ml⁻¹, 1 h) to eliminate contaminating bacterial nucleic acids, and phage DNA was subsequently purified using the DNeasy Blood & Tissue Kit (Qiagen) following the protocol beginning with proteinase K digestion. Sequencing libraries were prepared for Illumina sequencing, and reads were mapped to the *Escherichia* phage λ reference genome (GenBank accession number NC_001416) using Geneious Prime.

### Spot growth tests

*E. coli* strains harbouring the Retron-Sen3 system and derivatives or the corresponding control plasmids were grown overnight in LB supplemented with the appropriate antibiotics. Cultures were subjected to ten-fold serial dilutions, and 10 µl of each dilution were spotted onto LB agar plates containing the same antibiotics and, when required, 0.2% L-arabinose. The plates were then incubated overnight at 28 °C.

### ssDNA species isolation from immunoprecipitates and nuclease-based characterization

ssDNA species were isolated from FLAG immunoprecipitates as follows: 50 µl of immunoprecipitate was extracted with phenol:chloroform:isoamyl alcohol (25:24:1, pH 8), and nucleic acids were precipitated overnight at −80 °C with 2 volumes of 100% ethanol and 1/10 volume of 3 M sodium acetate (pH 5.2). The precipitated nucleic acids were collected by centrifugation at 14,000 rpm for 60 min at 4 °C. The pellets were air-dried and resuspended in 10 µL of distilled water. To detect ssDNA species, samples were first treated with RNase A/T1 (37 °C, 30 min) to remove RNA species and enrich the DNA component of ssDNA. To assess the chemical nature of the recovered product, aliquots were digested with 3′→5′ ssDNA Exonuclease I (ExoI) or 5′→3′ ssDNA exonuclease RecJ under the manufacturer’s recommended conditions. Sensitivity to both exonucleases confirmed that the recovered molecule corresponded to single-stranded DNA. For ssDNA species analysis, samples were mixed with TBE loading buffer, heated at 75 °C for 5 min, chilled on ice for 5 min, and resolved on a 10% denaturing polyacrylamide gel run in 1×TBE for 90 min at 20 W. The gels were stained for 15 min in a GelRed bath.

### Affinity purification of Retron-Sen3 RT–FLAG

*E. coli* strains harbouring plasmids expressing FLAG-tagged Retron-Sen3 RT or the corresponding untagged construct as a negative control were grown by inoculating 200 ml LB (with appropriate antibiotics) with 2 ml of an overnight culture. The cultures were incubated at 37 °C until an OD₆₀₀ of 1.2 was reached. Cells were harvested by centrifugation at 12,000 × *g* for 10 min at 4 °C, washed with 50 ml PBS, and resuspended in 8 ml lysis buffer (50 mM Tris–HCl pH 7.4, 150 mM NaCl, 1 mM EDTA, 1% Triton X-100) supplemented with protease inhibitors. Lysates were sonicated in four 15-s pulses and clarified by centrifugation at 12,000 × *g* for 10 min at 4 °C.

Anti-FLAG M2 agarose (100 µL; prepared according to the manufacturer’s instructions) was added to the cleared lysate, and the samples were incubated overnight at 4 °C with gentle rotation. The resin was collected by centrifugation at 8,200 × *g* for 30 s, and the supernatant was discarded. The beads were either used directly for co-incubation experiments (see below) or processed for the analysis of the Retron-Sen3 complex. For the latter, the resin was washed three times with 0.5 ml wash buffer (50 mM Tris–HCl pH 7.4, 150 mM NaCl). Bound proteins were eluted by incubating the resin with 100 µL wash buffer supplemented with 150 ng µL ⁻¹ 3×FLAG peptide for 30 min at 4 °C, followed by centrifugation at 8,200 × *g* for 30 s. The eluted material was stored at −20 °C.

### Proteomic analysis and differential enrichment analysis

Proteomic analysis was performed at the Proteomics Service of the Instituto de Parasitología y Biomedicina “López-Neyra” (CSIC, Granada). A total of 1.5 µg of protein from the affinity-purified sample was resolved for 10 min on a 4% SDS–PAGE gel and stained with Coomassie blue. The band of interest was excised and subjected to manual in-gel tryptic digestion for 18 h at 30 °C. Peptides were extracted using 0.2% trifluoroacetic acid (TFA) in 30% acetonitrile, dried, and stored at −20 °C until analysis. Prior to nLC–MS/MS, the peptides were resuspended in 0.1% formic acid (FA) and 0.3% acetonitrile. Samples were separated on an Easy nLC II system (Proxeon) coupled to an Amazon Speed ETD mass spectrometer (Bruker Daltonics, Bremen, Germany). Chromatographic separation was performed on a C18 reverse-phase analytical column (15 µm × 15 cm, 2.6 µm, 100 Å) using a 180-min gradient from 5% to 30% buffer B (buffer A: 0.1% FA in water; buffer B: 0.1% FA in acetonitrile) at a flow rate of 300 nl min⁻¹. The ion trap was operated in the m/z range of 390–1400. Database searches included carbamidomethylation of cysteine as a fixed modification and methionine oxidation as a variable one. Raw peptide-to-protein assignments were mapped to sample names, and emPAI values were organized into a protein × sample matrix for analysis. NA values (missing identifications) were replaced by 0, reflecting the absence of detection in label-free MS.

Differential protein abundance analysis was performed using LIMMA (Ritchie et al., 2015) in R (v4.3). For comparisons between C-terminally FLAG-tagged RT (RT–F) and untagged controls, a two-condition design matrix (“RT–F” and “Control”) was fitted using the lmFit function, followed by empirical Bayes moderation with `eBayes`. Log₂ fold changes and associated adjusted P-values were obtained using `topTable`. Comparisons between wild-type RT–F and the catalytic mutant (RTmut–F) were conducted analogously using a two-condition design (“RT–F” and “RTmut–F”). Proteins detected in at least one replicate of either condition were retained for analysis. For quantitative assessment of RT–HTH co-variation, exponentially modified Protein Abundance Index (emPAI) values were extracted for RT and HTH across all samples. Correlation analyses and scatter plots were generated in ggplot2 using `geom_point`, and linear regression was fitted using `geom_smooth (method = “lm”)`. The coefficient of determination (R²) and 95% confidence intervals were reported. Volcano plots were generated from LIMMA outputs using thresholds of |log₂FC| ≥ 1 and adjusted P < 0.05.

### Phylogenetic analyses

A maximum-likelihood tree was inferred from the conserved RT1–RT7 domains of 292 Type VI retrons in this study. The alignment was generated using MAFFT v7 (L-INS-i algorithm), and the tree was reconstructed with IQ-TREE2 using RTs from Types IV and V as outgroups to root the topology under the best-fitting model (F+I+R10) selected by ModelFinder using 1,000 SH-aLRT and 1,000 ultrafast bootstrap replicates to assess branch support. SP sequences associated with Type VI retrons were extracted from the RT–SP genomic contexts and assigned to subclades by phylogenetic analyses according to the classification of cognate RT. Sequences were aligned, and the alignment was manually curated to remove poorly informative positions, which is particularly important given the short length and high divergence of the SPs. Maximum-likelihood trees were inferred with IQ-TREE2 under the best-fitting model (LG+G4) using 2,000 SH-aLRT and 2,000 ultrafast bootstrap replicates to assess branch support.

### Structural modeling and alignment of SP toxins

Predicted 3D models of SP toxins were obtained from AlphaFold3 (Abramson et al., 2024) for *S. enterica* (Retron-Sen3 SP), *Y. pseudotuberculosis* (WP_038400095.1), *Xenorhabdus szentirmaii* (WP_038238213.1), and *V. parahaemolyticus* (Vpa2 SP). For each protein, the best-ranked model was selected and used for structural comparisons. Pairwise and multiple structural superpositions were performed using TM-align (Zhang and Skolnick, 2005) with default parameters to obtain TM-scores and aligned coordinate sets. Alignments were used to identify the conserved core domain, derive residue correspondence across orthologs, and generate the Sen3–Vpa2 structural alignment employed in subsequent analyses. Visualizations and downstream analyses were performed using Maestro 2024-3 (Schrödinger LLC, New York, NY) and ChimeraX (Meng et al., 2023).

### Hydropathy and hydrophobic moment analyses

Hydropathy and hydrophobic moment (μH) profiles were generated using custom R scripts. Hydropathy profiles were generated using the Kyte–Doolittle scale (Kyte and Doolittle, 1982) with a sliding window of seven residues, optimized for the short sequence segments analyzed. Hydrophobic moments (μH) were computed following the Eisenberg formalism (Eisenberg et al., 1984), applying a window size of 18 residues to capture the periodicity and overall amphipathic pattern characteristics of α-helical regions. All calculations, data handling, and plotting were performed in R (v4.3) using in-house scripts and the ggplot2 and dplyr packages.

### Domain mapping and schematic representation

Domain boundaries were defined from TM-align structural superpositions and supported by hydropathy (KD) and hydrophobic moment (μH) profiles of Sen3 and its SP orthologs. TM-align analysis identified an N-terminal extension in Sen3 SP, which was also present in the *Yersinia* and *Xenorhabdus* orthologs but absent in Vpa2. Each SP model was subdivided into three regions: the N-terminal segment, core domain, and C-terminal helix; based on conserved secondary structure organization and hydrophobic patterning. For Sen3 SP, the hydrophobic core and nucleus were defined as regions showing overlapping peaks of hydropathy and μH values. The amphipathic extension was defined as the C-terminal region with a hydrophobic moment vector greater than 0.35 Å. All domain schematics and comparative plots were generated in R (v4.3) using ggplot2, with Sen3 residue numbering as a reference. In Vpa2, the hydrophobic nucleus was annotated separately to indicate its distinct residue compositions.

## Results

### Genomic architecture and diversity of Type VI retrons

The Type VI retron group (Mestre et al., 2020), represented by 158 entries in our original dataset (RT-Clade 3), comprises systems with two principal genomic features. Approximately 78% encode a predicted Cro/C1-type HTH protein from cluster cl2_1, typically located upstream of or partially overlapping the RT gene. Within this group, approximately 27% additionally encode a small accessory protein (SP) from one of four conserved SP clusters, defining the canonical SP–HTH–RT organizational arrangement. Three SP clusters are equally represented among these systems: cl2075, cl2760, and cl3272, each accounting for one third of SP-containing entries, with cl3897 present as a minority lineage. No entry encodes more than one SP cluster simultaneously, consistent with strict modularity of the SP component within this retron type.

In the original bioinformatic characterization of Type VI retrons, no structured non-coding RNA (ncRNA) equivalent to the canonical msr–msd RNA was identified (Mestre et al., 2020). Independently, a systematic census of retrons across all 13 types found no detectable ssDNA species in any of the six Type VI representatives tested: *Pseudomonas* sp., *Bacteroides fragilis*, *Thiomonas delicata*, *Vibrio parahaemolyticus*, *Psychrobacter* sp., and *Candidatus* Levybacteria bacterium, corresponding to entries Mestre-604, -613, -662, -685, -703, and -729 in our original dataset, when expressed in *E. coli* (Khan et al., 2025). Zhang et al. (2025) subsequently retested Retron-Vpa2 (*V. parahaemolyticus*, Mestre-685) and Retron-Eco12, a phylogenetically related system, as complete operons under their native promoters; neither produced detectable ssDNA species under basal conditions, consistent with the results of Khan et al. (2025).

The absence of detectable ssDNA species across all experimentally tested Type VI retrons under standard expression conditions prompted us to examine additional members of the group more systematically. We focused on retrons encoding an SP from cluster cl2075, one of the three equally represented SP-associated lineages in Type VI. The cl2075 group forms a well-supported monophyletic RT subclade within Type VI (see below), predominantly associated with Beta- and Gammaproteobacteria including multiple Enterobacterales species, making these retrons particularly amenable to comparative genomic analysis and heterologous expression in *E. coli*.

To identify suitable candidates for functional characterization, we used a *Yersinia enterocolitica* retron sequence (Mestre-578) as a query to search for homologous systems within Enterobacterales. This search identified an orthologous retron in *Salmonella enterica*, encoding an SP from cl2075 (62% pairwise identity), an HTH protein from cl2_1 (79% pairwise identity), and an RT (63% pairwise identity) relative to the corresponding *Yersinia* components (**Figure 1A**). This retron (hereafter referred to as Retron-Sen3) was selected for detailed functional characterization.

**Figure 1.**
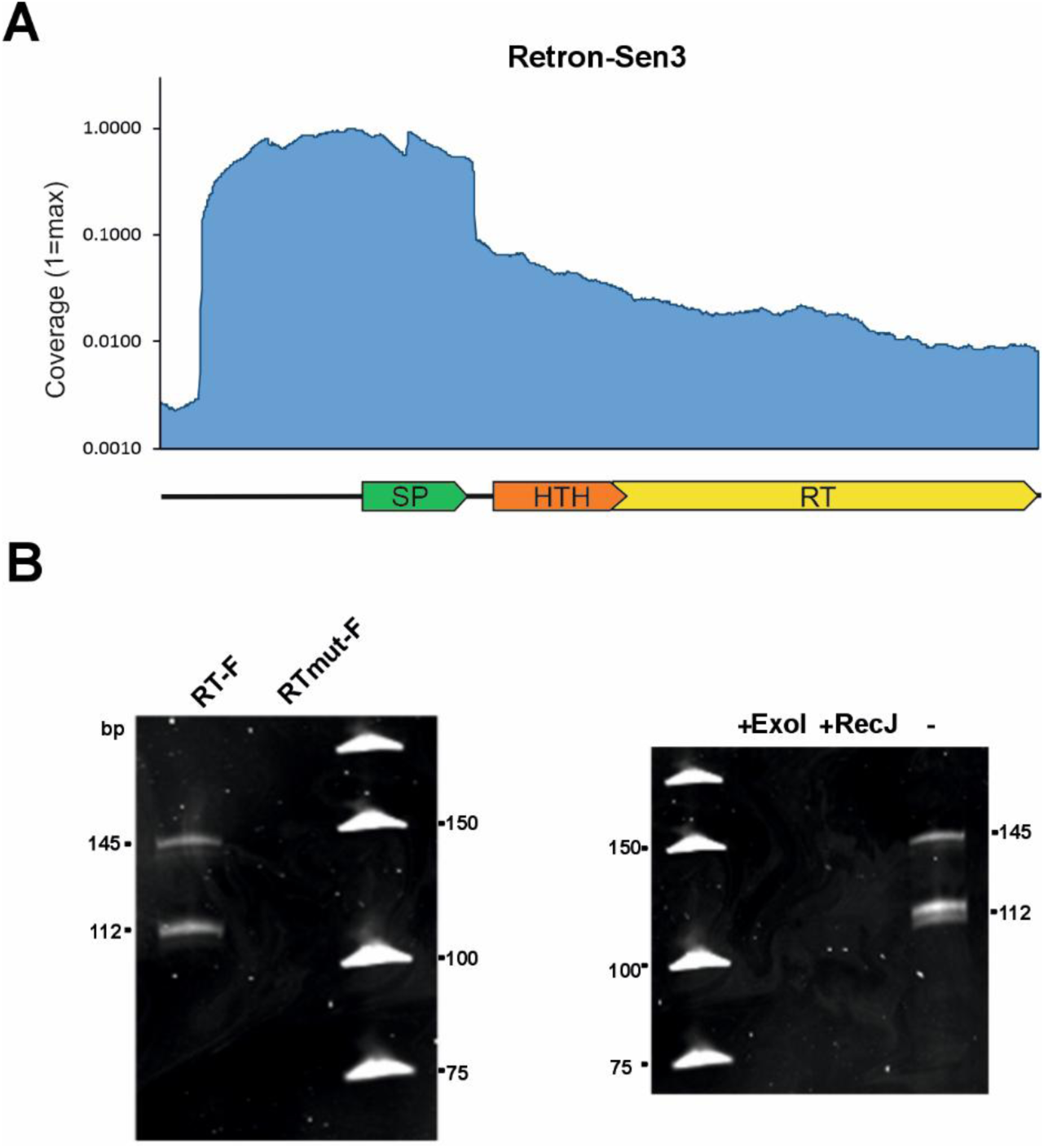
Genomic architecture, ssDNA species production, and associated RNA of Retron-Sen3. (**A**) Schematic representation of Retron-Sen3 genomic organization and coverage plot from RNA-seq of *E. coli* strain expressing the wild-type Retron-Sen3 module. (**B**) Denaturing PAGE analysis of nucleic acids extracted from anti-FLAG immunoprecipitates of *E. coli* cultures expressing Retron-Sen3 with a C-terminal 3×FLAG-tagged RT (RT-F). Samples were treated with RNase A/T1 prior to electrophoresis. An RT catalytic-site mutant (DD193–194AA)-FLAG (RTmut-F) was included as a negative control. Sensitivity to digestion by ExoI and RecJ confirms the single-stranded DNA nature of both products; an untreated sample (–) is shown for comparison.

### Retron-Sen3 produces RT-dependent ssDNA species and encodes a conserved retron-like ncRNA architecture

To determine whether Retron-Sen3 is capable of generating DNA species associated with retron activity, we expressed the complete SP–HTH–RT module under its native promoter in *E. coli*, using an RT fused to a C-terminal 3×FLAG tag (**Supplementary Figure S2**). Nucleic acids extracted from anti-FLAG immunoprecipitates and treated with RNase A/T1 yielded two discrete bands of approximately 112 and 145 nt; the smaller band consistently appeared as a closely spaced doublet, with one species more abundant than the other. Both products were absent in cells expressing a catalytic-site mutant (DD193–194→AA), establishing their dependence on RT activity, and both were sensitive to digestion by the 5′→3′ ssDNA exonuclease RecJ and the 3′→5′ ssDNA exonuclease ExoI (**Figure 1B**), confirming their single-stranded DNA nature and the accessibility of both termini. The direct sensitivity to RecJ (which requires a free 5′ end) indicates that these ssDNA species accumulate in a debranched form in *E. coli*, analogous to a subset of retrons in which the 2′–5′ RNA–DNA linkage is resolved *in vivo* by host nucleases prior to extraction (Khan et al., 2025). Together, these results establish that Retron-Sen3 produces discrete RT-dependent ssDNA species constitutively, at levels detectable by standard gel electrophoresis in the absence of phage infection, distinguishing it from all previously examined Type VI retrons (Khan et al., 2025; Zhang et al., 2025).

Given that retron ssDNA synthesis requires an RNA template, we next sought to identify the corresponding ncRNA of Retron-Sen3. RNA-seq performed on *E. coli* cells expressing the complete Retron-Sen3 module revealed a discrete enrichment of sequencing reads across the retron locus (**Figure 1A**). This enrichment spanned the region upstream of the conserved ncRNA element, the ncRNA itself, the SP coding sequence, and the downstream intergenic region extending to the ribosome-binding site of the HTH gene. This pattern was reproduced in cells expressing the RT catalytic mutant and an SP frameshift mutant (**Supplementary Figure S1**), confirming that transcript accumulation at this locus is independent of both RT catalytic activity and SP translation.

### Identification of a conserved ncRNA element across cl2075-associated Type VI retrons

To further characterize the RNA template associated with ssDNA synthesis and assess its conservation across the cl2075 lineage, we expanded the repertoire of Type VI retrons encoding RTs associated with SP from cluster cl2075. Using an HMM profile constructed from the RT sequences associated with cl2075 SPs in our original dataset (Mestre et al., 2020), we performed hmmsearch (HMMER v3.4) against an NCBI-compiled bacterial RT dataset comprising 26,799 non-redundant clusters derived from 107,067 sequences clustered at 90% identity. Phylogenetic analysis confirmed the membership of the retrieved sequences within the cl2075-associated group, yielding a total of 66 RTs and expanding the original collection approximately fivefold. For each RT, the 2-kb genomic regions flanking the coding sequence were extracted and screened for the canonical SP–HTH–RT architecture. Of the 66 retrieved RTs, 53 were confirmed to carry both a cl2075-associated SP and an HTH protein in the expected genomic arrangement and were used for subsequent comparative analyses.

To evaluate evolutionary constraints across this locus, we performed global multiple sequence alignments and quantified per-site conservation throughout the module. Conservation was unevenly distributed: the SP and HTH coding sequences were the most conserved (mean 0.62 and 0.61, respectively) and showed minimal length variability, consistent with strong purifying selection on both components. The RT gene exhibited intermediate conservation (mean 0.53) while maintaining a largely stable coding length, indicating preservation of core catalytic features alongside localized sequence diversification (**Figure 2A**). In contrast, the upstream region displayed lower overall conservation (mean 0.43) and substantially greater length heterogeneity. Rather than being uniformly divergent, this interval contained discrete local conservation maxima embedded within highly variable segments, including a particularly divergent internal stretch (mean conservation 0.33; mean non-gap length 92.6 nt, SD 23.6) (**Figure 2A**), suggestive of a structured element with conserved functional positions embedded in a variable scaffold.

**Figure 2.**
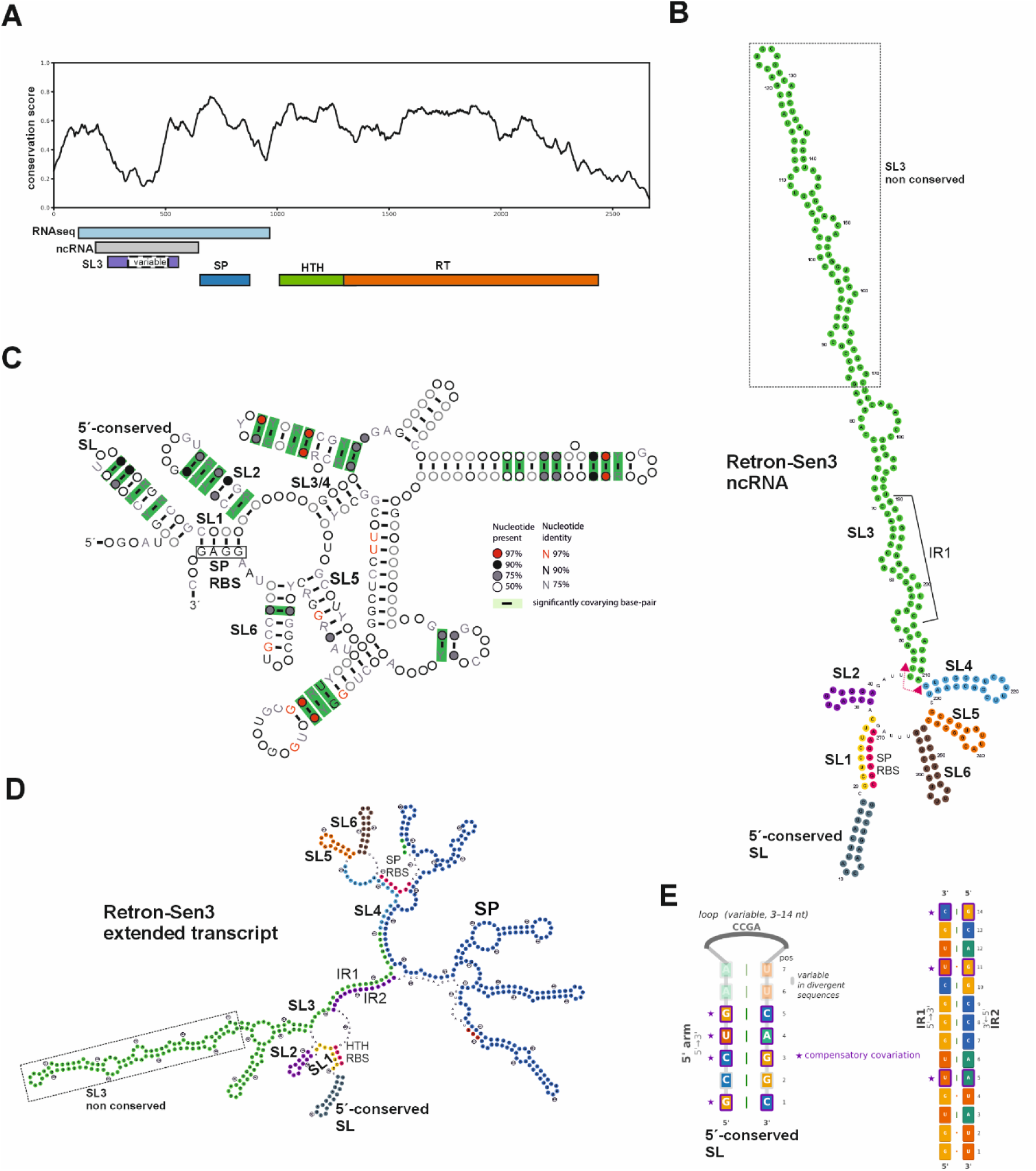
Evolutionary conservation and structural characterization of the cl2075-associated Type VI ncRNA. (**A**) Sliding-window conservation profile across the cl2075-associated Type VI retron locus. The RNA-seq-enriched interval, predicted ncRNA, SL3, and SL3 non-conserved variable region are indicated. Coding sequences corresponding to SP, HTH, and RT are annotated below. **(B)** Predicted secondary structure of the Retron-Sen3 ncRNA. Individual stem–loops (SL1–SL6) are shown in distinct colors, including the long internal SL3 (≈166 nt). The conserved stem–loop immediately upstream of the 5′ inverted repeat, the SL3 non-conserved variable region (dashed rectangle), the predicted SP ribosome-binding site (RBS), and the sequence within SL3 corresponding to IR1 are highlighted. The region deleted in the ΔncRNA mutant is indicated by a red double-headed arrow. (**C**) Consensus secondary structure of the cl2075-associated ncRNA derived from covariance analysis of 33 representative sequences, supported by 26 statistically significant covarying base pairs (observed-to-expected ratio = 1.9). The conserved architecture including SL1–SL6 is shown; the long internal domain corresponding to SL3 in Retron-Sen3 appears as a combined SL3/4 module in the consensus structure. Significantly covarying base pairs are highlighted in green. (**D**) Predicted secondary structure of the full-length Retron-Sen3 extended transcript encompassing the upstream ncRNA, the SP coding sequence, and the downstream intergenic region extending to the HTH ribosome-binding site. Individual stem–loops (SL1–SL6) are shown in distinct colors consistent with panel B. The 5′ (IR1) and 3′ (IR2) arms of the conserved distal inverted repeat flanking the SP coding sequence are indicated. The SP start codon (AUG, green), stop codon (UGA, red), SP coding sequence (blue), and ribosome-binding sites of SP and HTH (orange) are highlighted. Formation of the IR1–IR2 long-range pairing is predicted to involve a reorganization of the downstream stem–loops (SL4–SL6) within the extended structural framework. The SL3 non-conserved variable region is indicated (dashed rectangle). (**E**) Conservation of the stem–loop positioned immediately upstream of the 5′ inverted repeat across 53 cl2075-associated retrons (left) and detail of the IR1–IR2 long-range stem (right). The 5-bp stem of the upstream stem–loop is conserved across the dataset, with compensatory substitutions at positions 1, 3, 4 and 5; loop length varies between 3 and 14 nt. For the IR1–IR2 stem, compensatory substitutions are indicated at positions 5, 11 and 14. In both panels, nucleotides are colored by identity (G, yellow; A, teal; U, orange; C, blue), Watson-Crick pairs by vertical green bars, G:U wobble pairs by orange dots, and positions with compensatory substitutions by purple stars (★).

Motivated by the RNA-seq enrichment detected upstream of the SP gene in Retron-Sen3 and by this patterned conservation in the upstream interval, we examined this region in greater detail using structure-guided comparative analysis. Structure-guided alignments across the cl2075-associated retrons identified a conserved RNA element located within ∼280 nt of the SP start codon, displaying features consistent with a retron ncRNA: multiple stem-loops, discrete conservation peaks, and paired inverted repeats at the 5′ and 3′ termini (**Figure 2B**). A prominent internal stem–loop (designated SL3 in Retron-Sen3), spanning approximately 166 nt and containing a non-conserved central segment, is a candidate for the DNA stem–loop (DSL) generated during ssDNA synthesis. In Retron-Sen3, the predicted SP ribosome-binding site (RBS) is sequestered within the basal stem of the ncRNA structure, a feature also evident in the covariance model across the cl2075 lineage (**Figures 2B**). To assess whether this RNA architecture is conserved across the lineage, a covariance model (CM) derived from structure-guided alignments of 33 representative sequences was used to scan the corresponding genomic regions of all 53 cl2075-associated retrons, confirming the presence and boundaries of this structured element across the dataset, supported by 26 statistically significant covarying base pairs (observed-to-expected ratio = 1.9) in the consensus structure (**Figure 2C**). Notably, all members of this dataset (including Retron-Sen3) harbor a conserved stem–loop positioned immediately upstream of the 5′ inverted repeat, a feature not previously described in Type VI retrons **(Figure 2B)**.

### A conserved long-range RNA pairing links the ncRNA and SP coding regions across the cl2075 lineage

Having established the upstream ncRNA as a conserved structured element supported by covariance analysis, we next examined whether the extension of the transcript across the SP coding region and adjacent downstream region (**Figure 1A** and **Supplementary Figure S1**) reflects a conserved architectural feature of cl2075-associated retrons. The secondary structure analysis revealed the presence of an additional conserved inverted repeat flanking the SP gene: its 5′ arm (IR1) corresponds to a 3′ terminal segment of SL3, while the complementary arm (IR2) is located downstream of the SP stop codon. Formation of this long-range pairing would structurally integrate the upstream ncRNA domain with sequences flanking the SP coding region, suggesting that the SP gene is embedded within a broader RNA scaffold rather than constituting an isolated coding unit. In this alternative conformation, the SP ribosome-binding site, which is sequestered within the basal stem of the ncRNA structure (SL1) in the ncRNA conformation, is predicted to be repositioned into a stem formed with sequences corresponding to SL4, while the HTH ribosome-binding site is predicted to form a short stem with sequences immediately downstream of the 5′-conserved stem–loop (**Figure 2D**). The conservation of IR2 downstream of the SP stop codon, a region under no apparent coding or ncRNA structural constraint, combined with compensatory substitutions at three positions of the IR1–IR2 stem (positions 5, 11 and 14) that maintain base-pairing potential in RNA, is consistent with the structural integrity of this long-range interaction being preserved across the cl2075 lineage (**Figure 2E**).

To assess whether the region flanked by IR1 and IR2 is itself under structural selection, we performed covariance analysis on a structure-guided alignment of 52 cl2075-associated retrons spanning the IR1–IR2 interval. CaCoFold analysis identified 18 statistically significant covarying base pairs within the SP coding region (observed-to-expected ratio = 1.7; PPV = 100%), while the IR1–IR2 long-range stem itself showed no statistically significant covariation (**Supplementary Figure S3**). These results indicate that the SP coding sequence is itself under RNA structural selection, embedded within an extended transcript framework that represents a conserved architectural feature of cl2075-associated retrons.

### Evolutionary structure of Type VI retrons

Having established that Retron-Sen3 produces RT-dependent ssDNA species and encodes a conserved ncRNA, we sought to contextualize this system within the broader evolutionary diversity of Type VI retrons. Previous phylogenetic analyses of retrons have shown that Types IV, V, and VI form a monophyletic group (Mestre et al., 2020). To refine the evolutionary structure of Type VI retrons and determine the position of the cl2075-associated retrons analyzed here, we reconstructed the phylogeny of their RTs using a dataset that included all Type VI RTs from the original collection, the expanded set of cl2075-associated RTs, and the RTs of Types IV and V as outgroups. The resulting dataset comprised a total of 292 RT sequences. After rooting the tree with the Type IV and Type V sequences, Type VI RTs were recovered as a strongly supported monophyletic lineage (ultrafast bootstrap support = 100), clearly separated from both Type IV and Type V clades, which themselves formed a well-supported sister lineage (**Figure 3**).

**Figure 3.**
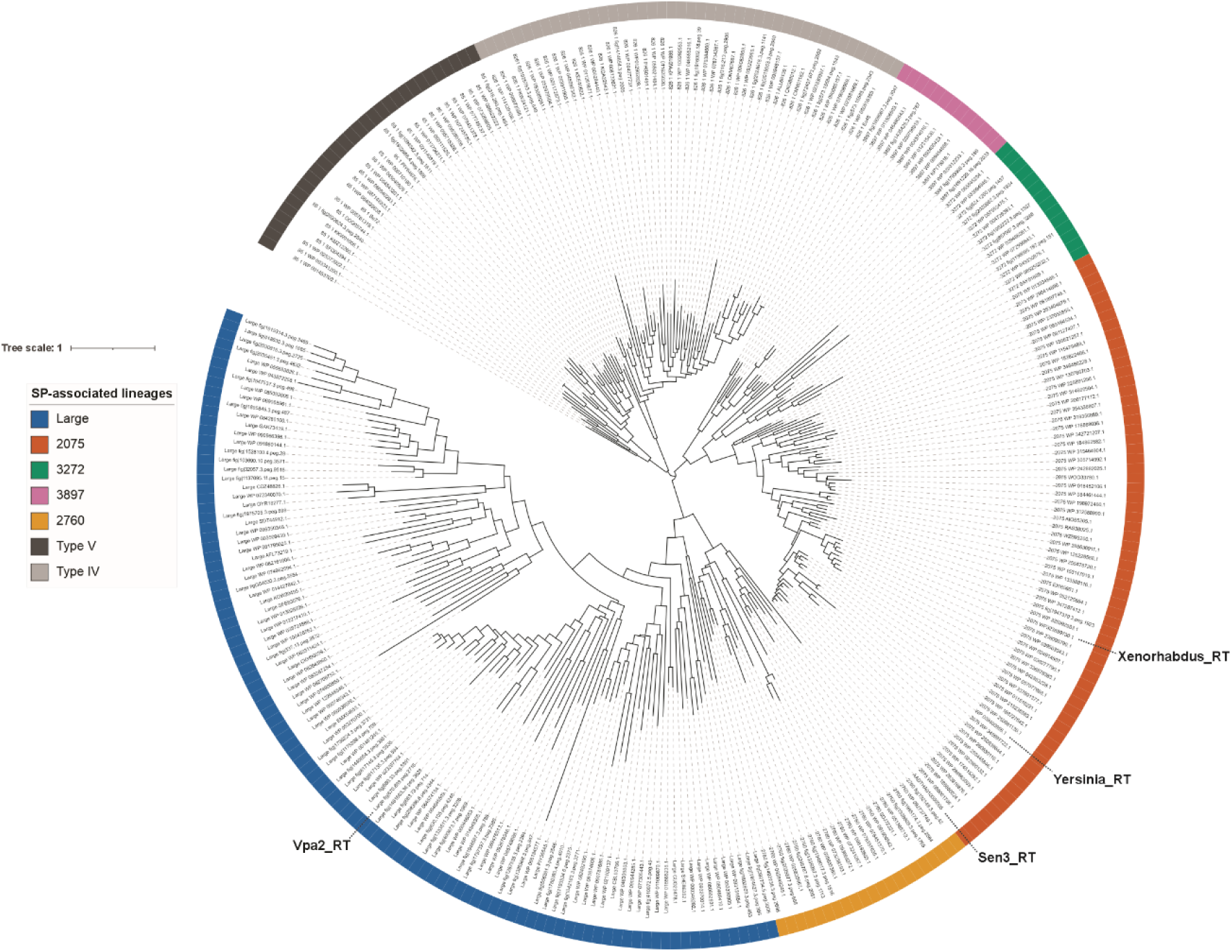
Phylogenetic structure of Type VI retrons. Maximum-likelihood phylogeny of reverse transcriptases (RTs) from Type VI retron systems reconstructed using IQ-TREE2 (Minh et al., 2020). Representative RTs from Type IV and Type V retrons were included as outgroups to root the topology. The resulting tree recovers Type VI RTs as a strongly supported monophyletic lineage distinct from Type IV and Type V. Within Type VI, five well-supported subclades are resolved, and clade nomenclature reflects the cluster identity of the associated small accessory protein (SP). Four subclades correspond to systems encoding SPs from clusters cl2075, cl2760, cl3272, and cl3897, whereas a heterogeneous lineage (Large) comprises RTs associated with SPs not belonging to these major clusters. The cl2075 subclade includes the expanded set of 66 representatives identified in this study. Two robust sister-group relationships are observed within Type VI: 2075 with 3272, and the “Large” lineage with 2760. The positions of Retron-Sen3, the *Y. enterocolitica* retron (Mestre-578), and the *Xenorhabdus szentirmaii* retron (Mestre-582) within the cl2075 subclade, and of Retron-Vpa2 within the “Large” lineage, are indicated. The tree was visualized using iTOL (Letunic and Bork, 2024). The corresponding Newick tree is included in **Supplementary Data 1**.

To examine diversification within Type VI, RT sequences were grouped according to the cluster identity of their associated SP proteins, and clade nomenclature refers to these SP clusters. Under this classification, the SP-associated groups 2075, 3272, 3897, and 2760 each formed well-supported monophyletic RT subclades (UFboot ≥ 97), indicating that RT diversification closely tracks SP cluster identity. In addition, a heterogeneous lineage (hereafter “Large”), comprising RTs associated with SP proteins that do not belong to any of the major SP clusters, also formed a strongly supported monophyletic clade (UFboot = 100). Within Type VI, the phylogeny reveals two robust sister-group relationships. The ‘Large’ and 2760 lineages form a maximally supported clade (UFboot = 100), indicating a shared evolutionary origin distinct from the remaining Type VI lineages. Likewise, the 2075 and 3272 groups constitute a well-supported sister pair (UFboot = 100), suggesting a second major lineage within the Type VI radiation. The 3897 clade emerges as an additional independent branch within Type VI, not forming a direct sister relationship with either of the two principal lineage pairs. The divergence patterns of these RTs correlate with host bacterial taxonomy. The 2075 and 3272 subclades are predominantly associated with members of the Beta- and Gammaproteobacteria, whereas the 2760 lineage is largely restricted to Bacteroidetes. The 3897 clade is primarily detected in *Pseudomonas* species, indicating a narrower host range. In contrast, the “Large” lineage exhibits broader taxonomic diversity across multiple bacterial groups consistent with a broader host range or more extensive horizontal dissemination.

Together, these results show that Type VI RTs constitute a cohesive evolutionary lineage structured into multiple internally well-supported sublineages, among which the cl2075 lineage represents a clearly resolved RT clade associated predominantly with Proteobacteria.

### HTH co-purifies with the Type VI retron RT

To assess whether HTH associates with the retron RT, we performed anti-FLAG immunoprecipitations of a C-terminally FLAG-tagged RT (RT-F), followed by label-free proteomics. Seven biological replicates expressing wild-type RT-F and three replicates expressing a catalytically inactive variant (RTmut-F) were analyzed. Comparison of RT-F samples with cells expressing untagged RT revealed consistent enrichment of the HTH protein in RT-FLAG immunoprecipitates (log₂ fold change ≈ 3; LIMMA analysis) (**Figure 4A**). To further evaluate the quantitative relationship between RT and HTH, we examined their co-variation across all samples. A scatter plot of emPAI values showed a clear positive correlation between RT abundance and HTH recovery across replicates (**Figure 4B**), consistent with stable association between both proteins.

**Figure 4.**
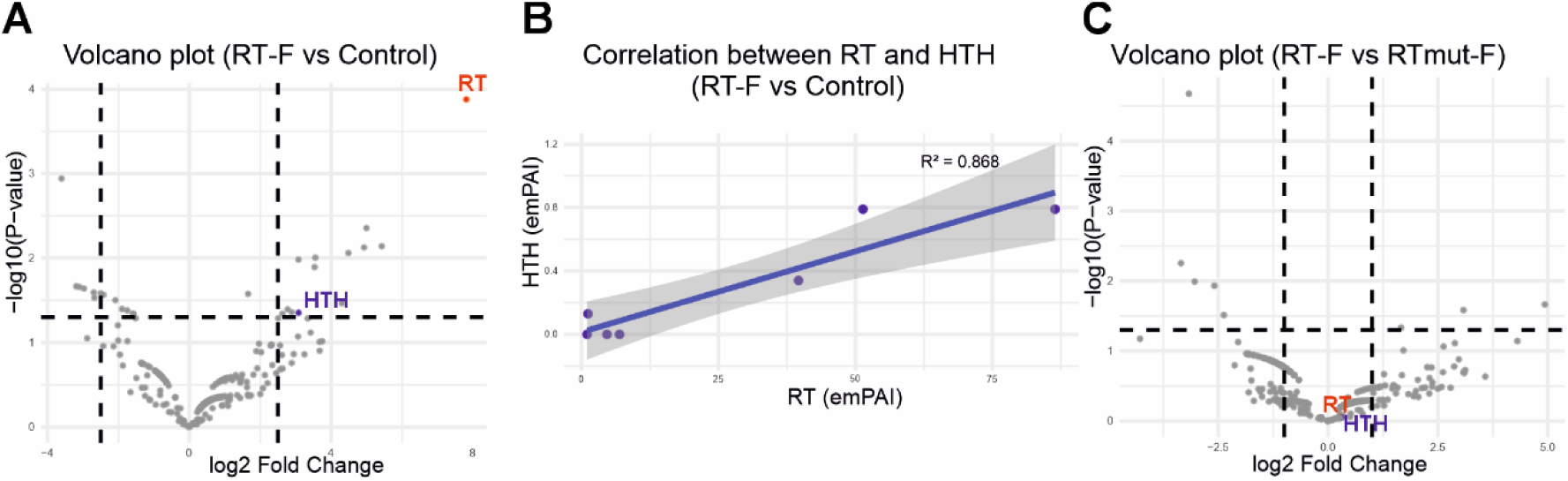
RT–HTH association is independent of RT catalytic activity. (**A**) Volcano plot comparing RT-FLAG (RT-F) immunoprecipitations with RT-untagged controls. The RT bait (red) and the HTH protein (blue) are enriched in RT-FLAG samples relative to controls. **(B)** Scatter plot showing the relationship between RT abundance (emPAI) and co-purified HTH across RT-F samples. A strong positive correlation is observed between RT and HTH levels. The fitted linear regression line is shown with its 95% confidence interval, and the corresponding coefficient of determination (R²) is indicated. **(C)** Volcano plot comparing RT-F and RTmut-F immunoprecipitations. RT and HTH display comparable abundance in wild-type and catalytic mutant samples, indicating that RT catalytic activity does not detectably influence RT–HTH association.

To determine whether reverse-transcriptase activity influences this interaction, we directly compared RTmut-F and wild-type RT-F samples (**Figure 4C**). Differential enrichment analysis revealed no significant differences in the abundance of either RT or HTH between catalytic mutant and wild-type conditions. Together, these data indicate that HTH reproducibly co-purifies with RT *in vivo* and that their association does not require RT catalytic activity. These findings support a functional interaction between HTH and RT within the Retron-Sen3 complex.

### Retron-Sen3 mediates phage defense through a toxin–antitoxin mechanism

We next tested whether Retron-Sen3 functions as an anti-phage defense system by expressing it in *E. coli* MG1655, a strain that does not encode a retron. Retron-expressing cells were challenged with a panel of 12 laboratory phages and 107 additional phages from the BASEL collection (Maffei et al., 2021; Humolli et al., 2025). Retron-Sen3 conferred robust resistance to phage λ, resistance to BAS01 and BAS81, and partial resistance to BAS23 and BAS41 (**Figure 5A**). The phage resistance phenotype was abolished by deletion of the predicted ncRNA SL3 region, frameshift mutation of the SP protein, or substitution of the RT catalytic residues, indicating that all three components are essential for defense. To further dissect the contribution of the ncRNA to retron-mediated defense, we examined the role of the conserved stem–loop located immediately upstream of the 5′ inverted repeat. Two variants were constructed: a deletion mutant lacking this stem–loop (Δ18nt-SL) and a replacement mutant in which the stem sequence was substituted with a non-pairing sequence predicted to disrupt the structure (18nt-SLmut). Both variants abolished Retron-Sen3–mediated resistance to bacteriophage λ, indicating that this conserved structural element is required for antiviral defense (**Figure 5A**). A variant carrying a frameshift mutation in the HTH protein could not be stably maintained in *E. coli*, indicating that HTH is required to maintain cell viability under basal conditions in the absence of phage infection, consistent with HTH acting as a negative regulator that suppresses toxicity associated with retron activity. Infection assays further revealed a multiplicity-of-infection (MOI)-dependent abortive infection phenotype (**Figure 5B**), consistent with defense activation leading to host cell death or growth arrest upon phage challenge.

**Figure 5.**
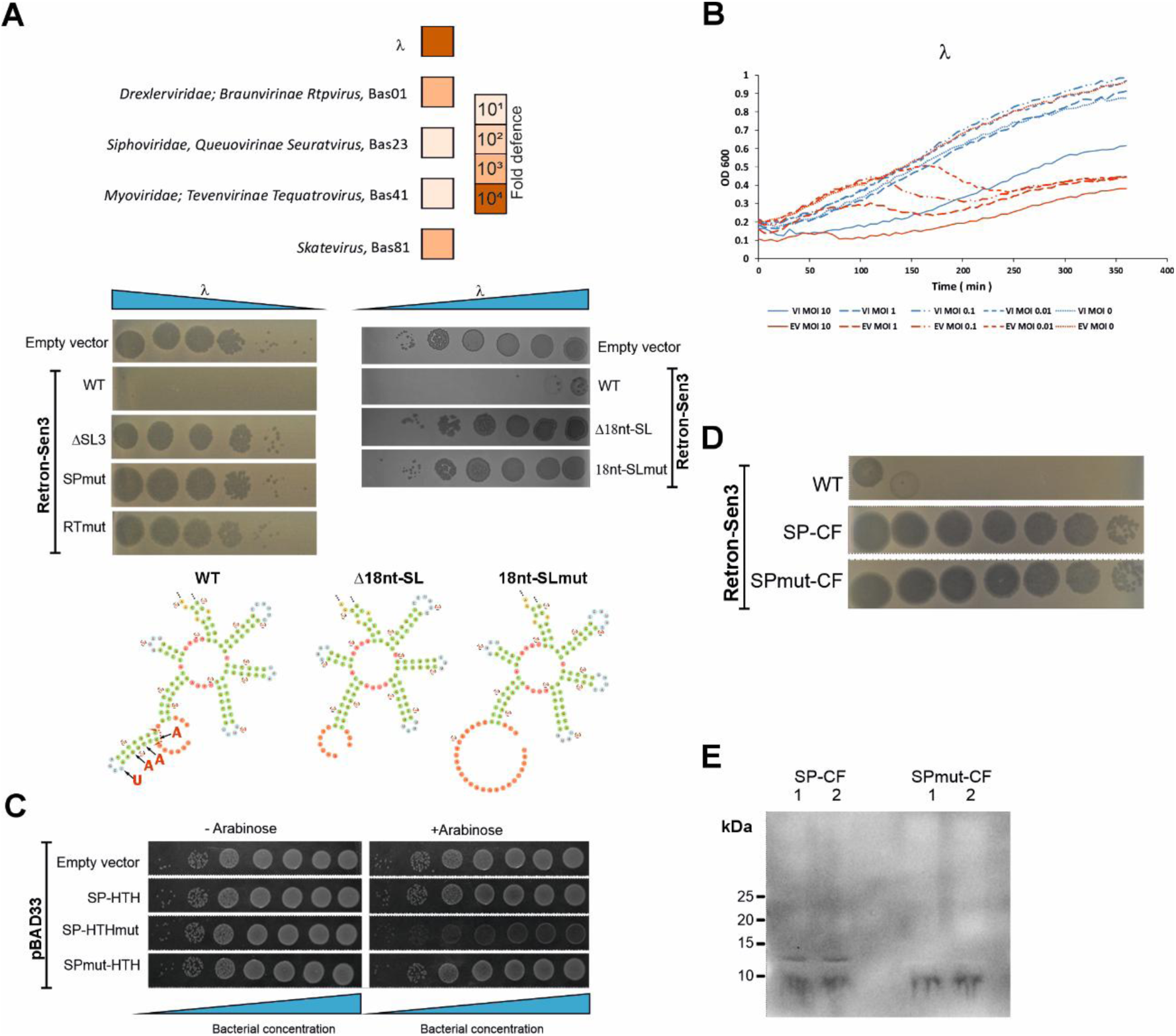
Retron-Sen3–mediated phage defense and functional dissection of its toxin–immunity module. **(A)** Fold defense conferred by Retron-Sen3 against susceptible bacteriophages. The complete Retron-Sen3 was expressed under its native promoter in *E. coli*. Strains carrying the empty vector, wild-type retron, deletion mutant lacking the SL3 stem–loop of the ncRNA, frameshift mutation in the SP toxin gene, or a catalytically inactive RT variant were challenged with bacteriophage λ. Robust resistance was observed only with the wild-type retron. The contribution of the conserved 18-nt stem–loop located immediately upstream of the ncRNA 5′ inverted repeat was further assessed using two additional variants: Δ18nt-SL, in which the stem–loop was deleted, and 18nt-SLmut, in which the stem sequence was replaced by a non-pairing sequence predicted to disrupt the structure. Schematics below the panels depict a simplified representation of the ncRNA region surrounding the 18-nt stem–loop for each construct. In the wild-type ncRNA, substituted nucleotides are indicated by arrows, whereas the deleted region is marked by a dashed line with arrowheads. The predicted structures of the mutant variants illustrate the loss of the conserved stem–loop. **(B)** MOI-dependent protection assay. Survival of strains expressing the wild-type retron was assessed across increasing multiplicities of infection (MOI 0.1–10). Retron-Sen3 provided dose-dependent protection that progressively diminished at high MOI, consistent with an abortive infection phenotype. **(C)** Minimal toxin–immunity module assay. The SP and HTH were expressed from an arabinose-inducible promoter either as a wild-type module or carrying inactivating mutations in SP or HTH. The wild-type SP–HTH module does not impair cell growth, and an SP loss-of-function variant was non-toxic. In contrast, inactivation of HTH resulted in marked toxicity, demonstrating that HTH is required to neutralize SP activity and establish self-immunity. **(D)** Plaque assay showing phage defense activity of Retron-Sen3 variants carrying FLAG-tagged SP proteins. Strains expressing SP with a C-terminal FLAG tag (SP-CF), an SP loss-of-function mutant with C-terminal FLAG tag (SPmut-CF) were challenged with bacteriophage λ. Defense activity was abolished in SP-CF, indicating that addition of a FLAG tag impairs SP function. (**E**) Western blot analysis confirming expression of the FLAG-tagged SP variants used in panel **D**. SP-CF is produced under basal conditions in the absence of phage infection. The SPmut-CF control confirms specificity of the FLAG signal. Two biological replicates are shown for each construct.

To further investigate the functional interplay between SP and HTH proteins, this region of the retron module, excluding the predicted ncRNA and RT, was expressed under an arabinose-inducible promoter. As shown in **Figure 5C**, co-expression of SP and HTH did not result in detectable toxicity, consistent with the phenotype observed for an SP loss-of-function mutant. In contrast, expression of SP in the presence of a non-functional HTH caused marked toxicity, indicating that SP is intrinsically toxic and that HTH counteracts this activity.

Expression of the full Retron-Sen3 carrying a C-terminal 3×FLAG-tagged SP protein abolished retron-mediated antiphage activity, despite detection of the tagged protein under basal conditions in the absence of phage infection (**Figure 5E**), suggesting that SP is produced constitutively and that its activity rather than its abundance is subject to regulation. The ability of HTH to suppress SP-mediated toxicity in the absence of the ncRNA and RT components suggests a direct functional interaction between SP and HTH.

### Retron-Sen3 recognizes phage λ through components of the Red operon

We next investigated how Retron-Sen3 recognizes and responds to phage infection by isolating phage mutants that escape retron-mediated defense. For phage λ, five independent escape mutants (λmut) were obtained, all of which were able to replicate efficiently in Retron-Sen3 expressing cells, although to different extents (**Figure 6A**). Genome sequencing revealed that all escaping phages carried mutations exclusively in the *exo* gene, which encodes the exonuclease component of the λ Red recombination operon, which also includes the *beta* and *gam* genes; Gam is a known inhibitor of RecBCD. These mutations consisted of single-nucleotide substitutions or frameshifts caused by nucleotide insertion (**Table S2**). Notably, ssDNA species production by Retron-Sen3 remained unchanged upon infection with either λ WT or λmut phages compared to uninfected cells (**Figure 6B**), indicating that activation of the defense system does not involve detectable modulation of ssDNA species production.

**Figure 6.**
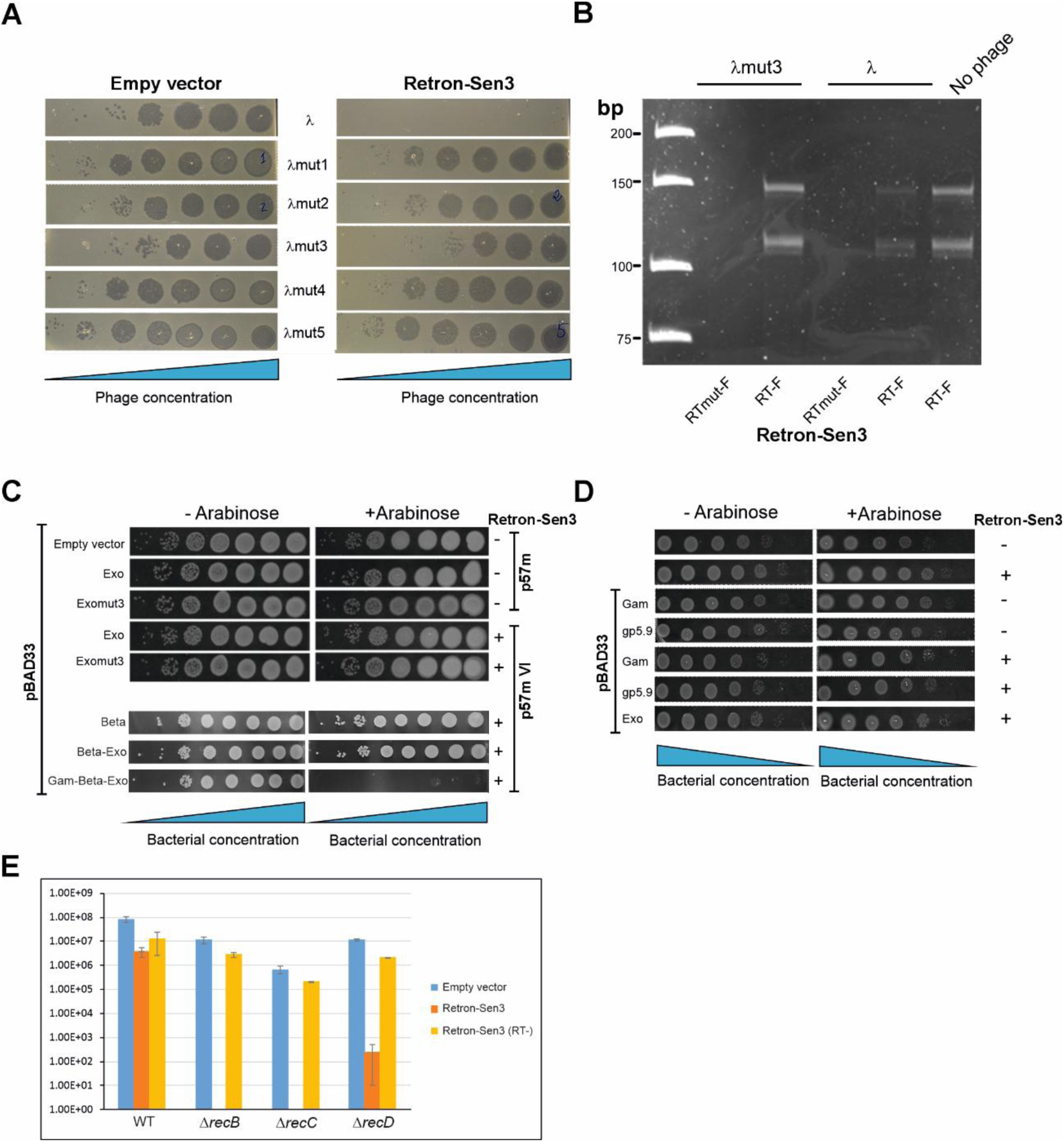
Interaction between the Retron-Sen3 defense system and phage λ. **(A)** Plaque assays comparing infection of *E. coli* strains carrying an empty vector or expressing Retron-Sen3 with wild-type phage λ and five independent escape mutants (λmut1–λmut5). Serial dilutions of phage stocks are shown from left to right (increasing phage concentration). Wild-type phage λ is efficiently blocked by Retron-Sen3, whereas all escape mutants bypass retron-mediated defense, although to different extents. **(B)** Denaturing PAGE analysis of ssDNA species produced by Retron-Sen3 during infection with wild-type phage λ or escape mutant λmut3, compared to an uninfected control. Nucleic acids were extracted from anti-FLAG immunoprecipitates of cells expressing RT-F or RTmut-F and treated with RNase A/T1 prior to electrophoresis. ssDNA species levels remain unchanged upon infection with either wild-type or escape mutant phage. **(C)** Effect of individual λ Red recombination proteins on Retron-Sen3 activation. The Retron-Sen3 module was expressed from its native promoter in plasmid p57m VI. Exo (wild type), Exomut3, Beta, Beta+Exo, Gam and the complete Red operon (Gam+Beta+Exo) were expressed from the arabinose-inducible vector pBAD33Only expression of the complete Red operon triggered Retron-Sen3–mediated defense, as evidenced by growth inhibition upon arabinose induction. **(D)** Same experimental setup as in panel **C**, testing individual proteins known to inhibit or interact with the host RecBCD complex. Gam from the λ Red operon, gp5.9 (a functional RecBCD inhibitor from phage T7), and Exo were expressed individually from pBAD33 in cells carrying Retron-Sen3. Serial dilutions of bacterial cultures are shown from left to right (increasing bacterial concentration). None of the individual proteins triggered Retron-Sen3–mediated defense. (**E**) Transformation efficiency of *E. coli* strains carrying mutations in the RecBCD pathway. Wild-type, Δ*recB*, Δ*recC*, and Δ*recD* strains were transformed with plasmids carrying an empty vector, wild-type Retron-Sen3, or a Retron-Sen3 variant carrying a catalytic-site mutation in the RT (RT-). Transformation efficiency is expressed as colony-forming units per microgram of DNA relative to the empty vector control. Δ*recB* and Δ*recC* strains show severely reduced transformation efficiency with wild-type Retron-Sen3, whereas the RT catalytic mutant restores transformability, indicating that retron-mediated toxicity is responsible for the observed growth inhibition. The partial tolerance observed in the Δ*rec*D background is also indicated.

To determine whether λ activates Retron-Sen3 through Exo or other components of the Red operon (Beta and Gam), we co-expressed the Retron-Sen3 module under its native promoter together with the corresponding phage genes placed under an arabinose-inducible promoter (**Figure 6C–D**). Expression of λ recombination proteins: Exo (wild-type or mutant), Beta, Beta together with Exo, or Gam alone, did not trigger Retron-Sen3 activation. In contrast, co-expression of the complete Red operon (Gam, Beta, and Exo) induced Retron-Sen3–mediated defense. These observations indicate that activation of Retron-Sen3 does not depend on a single Red protein but instead requires the coordinated activity of the Red recombination machinery.

*E. coli* cells carrying deletions of *recB* or *recC* could not be transformed with plasmid constructs harboring Retron-Sen3, whereas a Δ*recD* background still showed low but detectable transformation efficiency (**Figure 6E**). This inability to be transformed was overcome by introducing an RT catalytic-site mutation into the retron, indicating that the observed toxicity depends on retron activity. The partial tolerance observed in the ΔrecD background further suggests that retron activation correlates with impairment of RecBCD function. Together, these findings support a model in which Retron-Sen3 monitors RecBCD activity and becomes activated when this pathway is inhibited.

### Conserved hydrophobic core and divergent amphipathic organization in SP toxin C-terminal domains

Given that retron-mediated defense ultimately depends on the activity of the SP toxin, we next examined the structural organization of SP proteins associated with Type VI retrons. Previous bioinformatic analyses identified these proteins as small predicted helical proteins (Mestre et al., 2020). To evaluate structural relationships within cluster 2075, we selected representative orthologs that capture the two structural patterns observed in this group: the *Salmonella enterica* (Retron-Sen3) SP, which exhibits a two-helix topology, and the orthologs from *Yersinia pseudotuberculosis* (RT:WP_038400095.1) and *Xenorhabdus szentirmaii* (RT:WP_038238213.1), in which the long C-terminal helix of Sen3 is split into two shorter helices, producing a three-helix arrangement. Structural models predicted with AlphaFold3 (Abramson et al., 2024) were aligned using TM-align. The superposition (average TM-score ≈ 0.58; RMSD ≈ 2.1 Å over 53 aligned residues) delineates a conserved structural core formed by two N-terminal α-helices, whereas the extended C-terminal helix of the Sen3 toxin (∼residues 45–79) lies only partially within the superposition region. These results indicate that the proteins are structural homologs with a shared helical core but partially divergent C-terminal topologies (**Figure 7A**).

**Figure 7.**
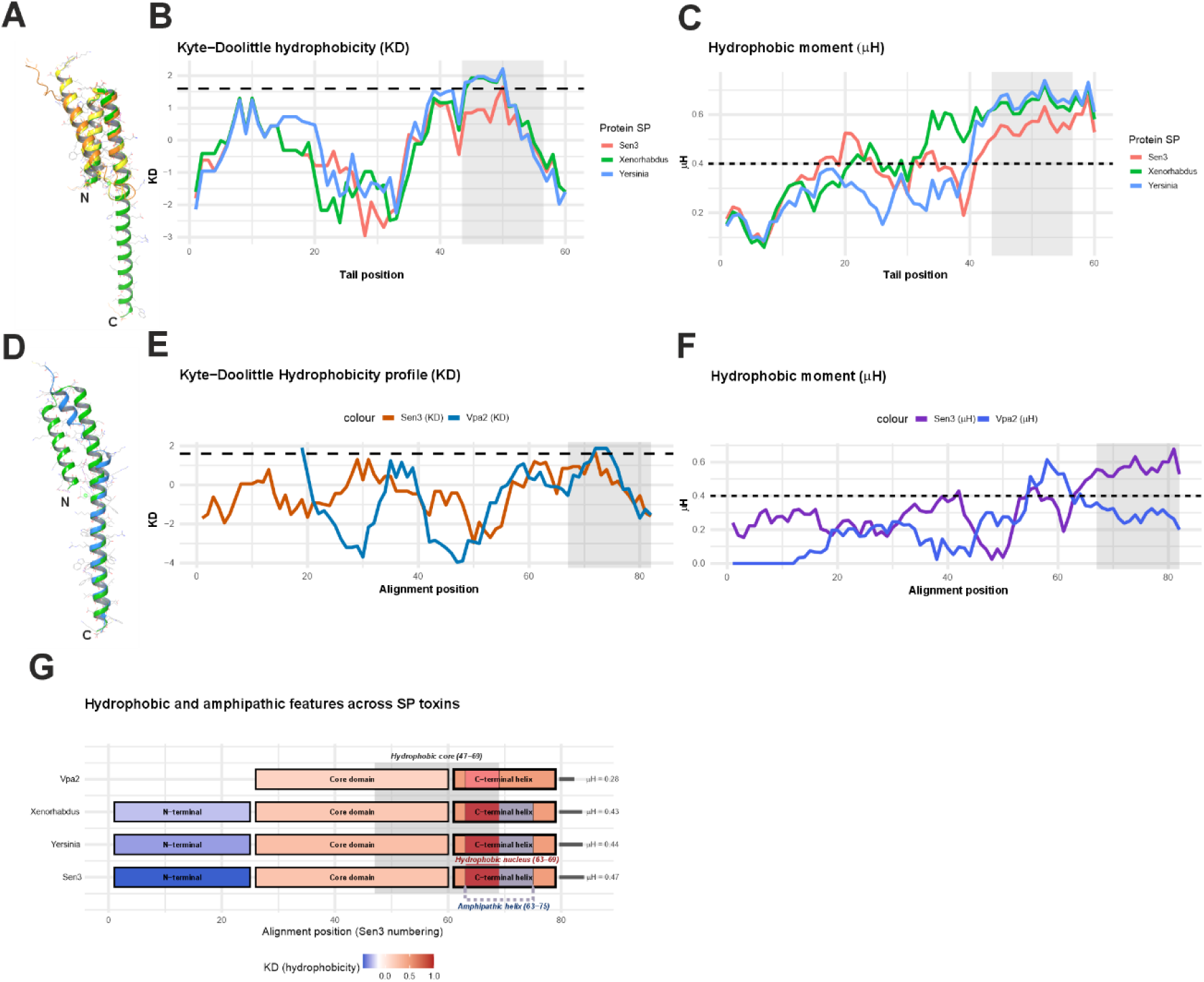
Comparative structural and biophysical features of Type VI retron-associated SP toxins. **(A)** TM-align superposition of the 3D models of SP orthologs from *S. enterica* (Sen3, green), *X. szentirmaii* (yellow), and *Y. pseudotuberculosis* (orange), showing conservation of the core helical topology despite differences in the organization of the C-terminal helices. **(B)** Kyte–Doolittle (KD) hydropathy profiles calculated across the C-terminal 60 residues of the SPs from *S. enterica* (Sen3), *X. szentirmaii* and *Y. pseudotuberculosis.* **(C)** Hydrophobic moment (μH) profiles for the same region assuming α-helical periodicity. The shaded region indicates the predicted C-terminal amphipathic α-helix in Sen3 (approximately residues 63–75), corresponding to the C-terminal helical segment observed in the structural models. **(D)** TM-align structural superposition of Sen3 SP (green) and the phylogenetically distant Vpa2 SP from *V. parahaemolyticus* (light blue), illustrating conservation of the core fold despite divergence in N-terminal architecture. **(E)** Kyte–Doolittle hydropathy profiles mapped onto the structural alignment of the core region defined by TM-align between Sen3 and Vpa2. **(F)** Corresponding hydrophobic moment (μH) profiles for the same aligned region, highlighting reduced amphipathic polarity in Vpa2 helix compared to Sen3. In panels B–C the x-axis corresponds to the C-terminal 60 residues of the SP proteins, whereas in panels E–F positions correspond to the TM-align structural alignment. **(G)** Schematic representation of the domain organization of Sen3 SP and its orthologs compared with Vpa2. All coordinates are indicated according to the Sen3 SP reference sequence. The C-terminal region contains an amphipathic α-helix (residues 63–75) that includes a hydrophobic nucleus (63–69), highlighted within a broader hydrophobic core (47–69) defined from Kyte–Doolittle profiles. The amphipathic helices of Sen3, Yersinia and Xenorhabdus are annotated with their corresponding hydrophobic moment (μH) values. In Vpa2, a truncated C-terminal helix retains a weak hydrophobic nucleus aligned to residues 63–69 of Sen3 but lacks the amphipathic extension (**Supplementary Table S3**).

To investigate whether this structural divergence affects the physicochemical organization of the C-terminal region, we analyzed the hydropathy and hydrophobic moment (μH) profiles of the last 60 residues of each ortholog (**Figure 7B-C**). The Kyte–Doolittle plots revealed a conserved hydrophobic segment between residues ∼45 and 55, where all three proteins display pronounced local maxima (KD ≈ 1.5–2.0). Consistent with this pattern, μH profiles calculated assuming an α-helical configuration showed coincident peaks in the same region (μH ≈ 0.5–0.6), indicating the presence of a conserved amphipathic helical segment at the extreme C-terminus. Although this region is not fully captured in the TM-align superpositions owing to helix-splitting events in the *Yersinia* and *Xenorhabdus* orthologs, the hydropathy and μH analyses demonstrate that its underlying physicochemical signature is conserved across cluster 2075.

To assess structural divergence beyond cluster 2075, we compared the Sen3 SP toxin with the phylogenetically distant SP protein encoded by Vibrio parahaemolyticus Vpa2, a representative of the “Large” Type VI retron clade. TM-align superposition of the 79-residue Sen3 SP with the 55-residue Vpa2 model revealed a common structural core spanning residues S22–K79 of Sen3, yielding an RMSD of 2.43 Å over the aligned residues and a TM-score of 0.70 (normalized to the shorter protein). Despite negligible sequence identity (1.9%), the TM-score of 0.70 indicates significant structural similarity (**Figure 7D**). Notably, Sen3 and other cl2075 SP proteins contain an N-terminal extension of approximately 25 residues that lies outside this structural core, whereas Vpa2 lacks this extension entirely. This observation suggests that the aligned region represents the evolutionary core of the SP fold, while the N-terminal extension found in cl2075 members may contribute to lineage-specific functional interactions.

To determine whether structural similarity between Sen3 and Vpa2 extends to their surface physicochemical properties, we compared hydropathy and μH profiles across the aligned C-terminal region (Sen3 residues 22–79). Kyte–Doolittle plots revealed broadly comparable hydrophobicity patterns, with alternating hydrophobic and hydrophilic segments and a pronounced hydrophobic peak near the C-terminus (**Figure 7E**). This peak corresponds to the conserved hydrophobic segment previously identified among cl2075 orthologs, reinforcing its role as a common feature of the SP toxin family.

In contrast, μH profiles revealed a clear divergence in amphipathic organization (**Figure 7F**). In Sen3, μH values progressively increase toward the C-terminal tail and reach a pronounced maximum (μH ≈ 0.6), consistent with a well-defined amphipathic α-helix. In Vpa2, μH values peak earlier and decline toward the extreme C-terminus (μH ≈ 0.3–0.4), indicating a reduced amphipathic character in the corresponding region. Thus, although both proteins share a conserved C-terminal hydrophobic core, only the cl2075 SP toxins (including Sen3) display the amphipathic polarity characteristic of the extended C-terminal helix. In Vpa2, the corresponding region is shorter and enriched in aromatic residues, resulting in a more uniformly hydrophobic segment. The pronounced amphipathic character of this helix in cl2075 SP toxins suggests that it may mediate specific protein-protein interactions or other functional properties that are absent in Vpa2. Together, these results indicate that SP toxins share a conserved structural nucleus but differ in the organization of their terminal helices, revealing lineage-specific adaptations within the Type VI retron family (**Figure 7G** and **Table S3**).

### Structural modelling of Sen3 HTH–SP and RT–HTH–SP complexes

To explore a possible structural basis for the functional interaction between SP and HTH suggested by the co-expression toxicity assays described above, we modelled the HTH–SP complex using AlphaFold3(Abramson et al., 2024). In the predicted models, the dominant interface is constituted by the α3-helix and C-terminal region of the SP (residues 53–79) and the α6-helix of the HTH (residues 76–91) (**Figure 8A-C**), matching the hydrophobic and amphipathic signatures identified by Kyte–Doolittle and hydrophobic moment (μH) analysis. The predicted interface residues (54–55, 57–58, 60–65, 68–74 and 77–79) map onto this surface, consistent with a mode of interaction involving the hydrophobic face of this helix (**Figure 8B**). A secondary N-terminal (Nt) contact patch from α1-helix of SP (residues 1–23) and the α6-helix of the HTH was also predicted, likely providing stabilizing rather than determinant interactions (**Figure 8A-C**). The models further suggest that Sen3 SP may display local structural plasticity upon HTH engagement: whereas the isolated SP adopts a two-helix bundle, the complex stabilizes a short intermediate helix, yielding a three-helix arrangement. In contrast, SP orthologs from *Yersinia* and *Xenorhabdus* intrinsically favor a three-helix topology, suggesting local plasticity in the SP fold and HTH-dependent stabilization of this intermediate segment.

**Figure 8.**
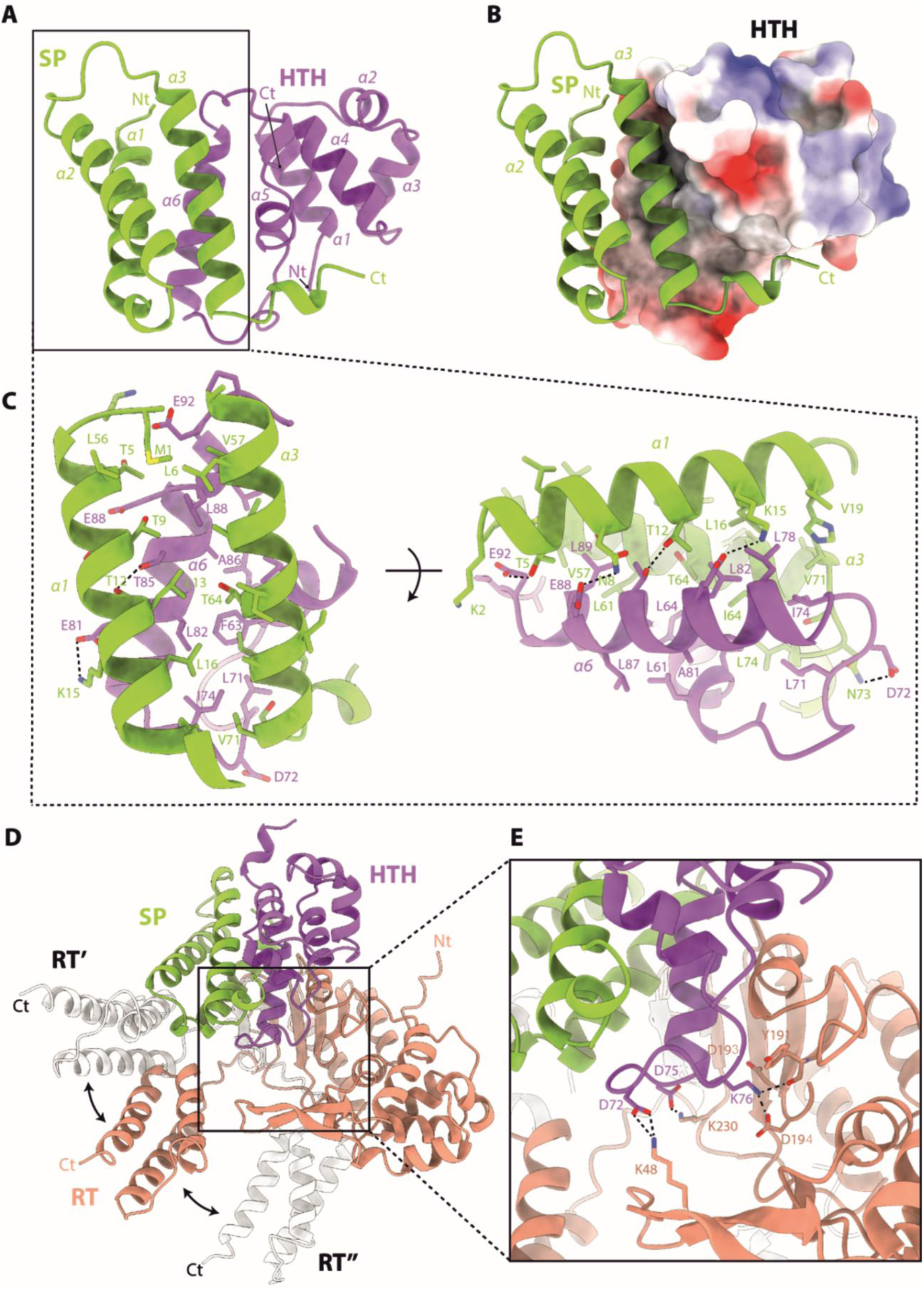
Structural modelling of the Sen3 HTH–SP and RT–HTH–SP complexes. **(A)** Structural prediction of the SP-HTH binary complex in cartoon representation. SP protein is displayed in green while HTH protein is depicted in purple. **(B)** Electrostatic potential surface of HTH in complex with SP (cartoon), showing the hydrophobicity of the α6 helix from HTH. The interaction is dominated by the interaction of α6 helix from HTH with helices α1 (residues 1-23) and α3 (residues 53-79) from SP. **(C)** Zoom view of the SP-HTH interface displaying the key interactions between both proteins. **(D)** Structural prediction of the SP-HTH-RT ternary complex. RT protein is depicted in Salmon, while SP and HTH proteins follow the same color code in panel **A. (E)** Close-up view of the RT catalytic center showing the proximity between the HTH and the YVDD (191–194) motif. This motif and the HTH-RT key interacting residues are displayed in sticks.

To explore how the toxin–antitoxin module integrates with reverse transcriptase, we modelled the RT–HTH–SP ternary complex by using Alphafold3 (Abramson et al., 2024). All models generated converged, except for the C-terminal (Ct) region spanning residues 236 to 319 (**Figure 8D**). Although the substrate is absent in the models, the predicted geometry convergently positions the HTH adjacent to the conserved YVDD motif (Y191–D194) of the RT. Within this region, there are three salt bridge interactions between K76, D72 and D75 from HTH with D193, K48 and K230 from RT, respectively. In addition, K76 of the HTH lies within the hydrogen-bonding distance of the Y191 main chain carboxyl group (**Figure 8E**). However, biochemical analyses indicate that HTH binding does not impair RT catalytic activity (**Figure 4C**), suggesting that the RT–HTH interaction serves a regulatory rather than catalytic role. Together, these models outline a structural framework in which HTH establishes coordinated interfaces with both SP and RT, consistent with its proposed function as an antitoxin that integrates information from the RT–ncRNA complex to regulate SP activity.

## Discussion

Type VI retrons have recently emerged as mechanistically unusual members of the retron superfamily. Whereas previously characterized retrons commonly rely on msDNA-dependent control of toxic effectors, recent work on Retron-Vpa2 suggested a distinct strategy in which msDNA promotes toxin translation during phage infection (Zhang et al., 2025). Our characterization of Retron-Sen3 now reveals a second and fundamentally different solution within Type VI retrons. In this system, RT-dependent ssDNA species are produced constitutively, SP toxin is present under basal conditions, and defense is controlled at the post-translational level through HTH-mediated toxin restraint. These findings indicate that Type VI retrons are not governed by a single regulatory mechanism, but instead comprise multiple lineage-specific strategies for coupling reverse transcription to phage defense.

Retron-Sen3 appears to exist in a constitutively primed defensive state. RT-dependent ssDNA species are maintained under basal conditions and remain unchanged during infection, while the intrinsically toxic SP protein is present but restrained by HTH, which is required for basal viability. These observations indicate that regulation occurs primarily at the level of toxin activity rather than toxin abundance. In parallel, comparative analyses indicate that the SP coding region is embedded within a conserved structured RNA scaffold, suggesting that RNA architecture remains an important organizing feature even in this post-translational regulatory system. This organization contrasts with the inducible translational control described for Retron-Vpa2 and indicates that activation in Sen3 occurs downstream of ssDNA synthesis and toxin production.

Phage genetics and host mutational analyses indicate that inhibition of the RecBCD pathway is the proximal trigger for Retron-Sen3 activation. All λ escape mutants mapped to *exo* within the Red recombination operon, and activation required the complete Red module, Gam, Beta, and Exo acting together, whereas individual components or Beta+Exo were insufficient. This contrasts with Retron-Vpa2, in which Beta+Exo alone is sufficient to trigger defense (Zhang et al., 2025), indicating a mechanistically distinct activation pathway in Sen3. The essential requirement for Gam, a known inhibitor of RecBCD, together with the toxicity observed in *recB* and *recC* mutant hosts, supports a surveillance model in which Retron-Sen3 monitors RecBCD activity and becomes activated when this pathway is inhibited during phage infection. Notably, ssDNA species levels remained unchanged upon infection, further confirming that activation occurs independently of detectable changes in ssDNA abundance.

Multiple lines of evidence identify HTH as the central regulatory node of the Sen3 system. HTH suppresses SP toxicity independently of RT and ncRNA, yet also associates with RT irrespective of catalytic activity, indicating that its interactions with the toxin and reverse-transcription modules are functionally separable. This dual connectivity places HTH at the interface between constitutive ssDNA biogenesis and toxin control, providing a plausible mechanism by which the state of the retron complex could be coupled to defense activation. Structural models of the RT–HTH–SP ternary complex are consistent with this organization, placing HTH adjacent to the RT catalytic cleft without impairing enzymatic activity, although the biochemical basis of these interactions remains to be established.

The placement of Sen3 within the conserved cl2075 lineage suggests that this post-translational regulatory mechanism may extend beyond a single retron system. Structural comparisons further indicate that SP toxins retain a conserved two-helix core despite substantial sequence divergence across Type VI lineages. In Sen3, a distinctive amphipathic C-terminal extension is predicted to form the principal interface with HTH, providing a possible structural basis for toxin neutralization and reinforcing the proposed connector role of HTH within the ternary complex. The absence of this extension in the phylogenetically distant Vpa2 toxin highlights the distinct interaction topologies that may underlie mechanistic divergence among SP families.

Together, our findings establish Retron-Sen3 as a constitutively primed Type VI defense system controlled through a post-translational toxin–antitoxin mechanism. In this architecture, constitutive ssDNA biogenesis coexists with basal toxin production, while HTH-mediated restraint prevents self-toxicity until phage-induced perturbation of RecBCD is sufficient to trigger activation. More broadly, these results demonstrate that shared genomic architecture within Type VI retrons masks substantial mechanistic divergence, with distinct lineages repeatedly repurposing reverse transcription, structured RNAs, and toxic effectors into alternative antiviral control circuits.

## Acknowledgments

We thank Ascensión Martos Tejera and José María del Arco Martín for technical assistance.

## Author contributions

F.M.G-R performed the experimental work and contributed to writing, reviewing, and editing the manuscript. F.M.A. contributed to the identification and selection of the *S. enterica* Retron-Sen3, and reviewing the manuscript. R.M. contributed to the structural modeling, writing and reviewing the manuscript. N.T. supervised the study, conducted the ncRNA identification, structural and phylogenetic analyses, wrote the manuscript, and contribute to reviewing, and editing.

## Supplementary Figures

**Supplementary Figure S1.**
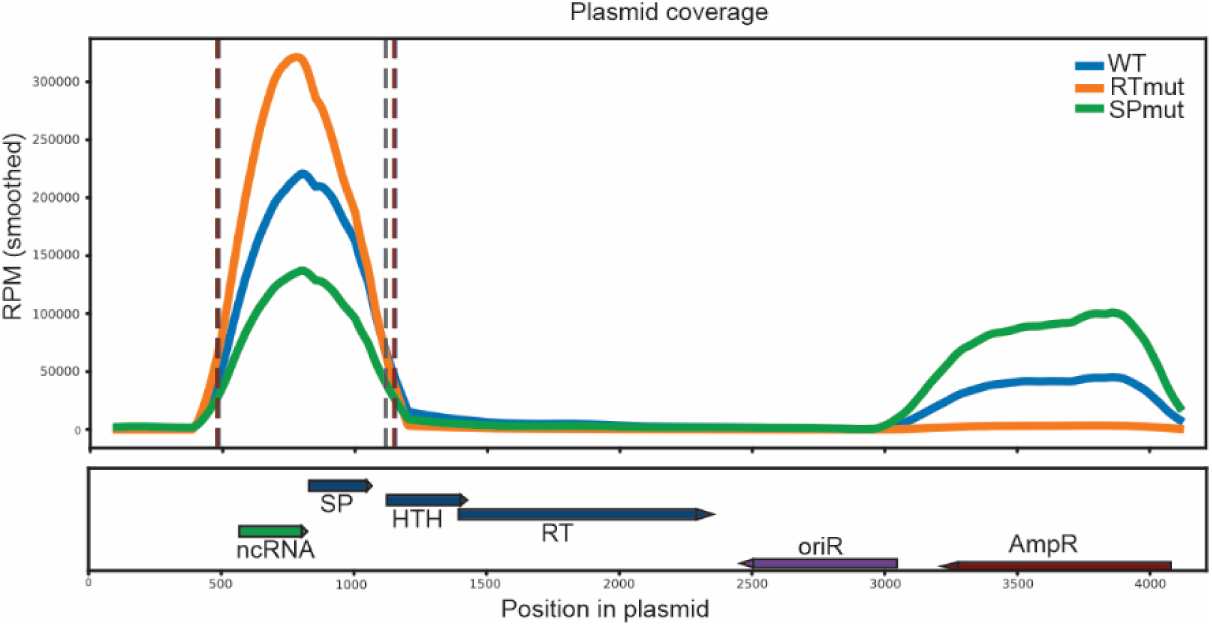
RNA-seq coverage across the Retron-Sen3-containing plasmid. Distribution of RNA-seq reads mapped to the plasmid sequence in strains expressing the Retron-Sen3 module, including WT, the RT catalytic mutant (RTmut), and an SP frameshift mutant (SPmut). Coverage profiles are shown across the entire plasmid to illustrate the global distribution of reads. Coverage values are expressed as reads per million (RPM) and normalized to the total number of reads mapped to the plasmid in each sample. Coverage traces were smoothed using a rolling average (200-nt window) to facilitate visualization. Vertical dashed lines indicate the boundaries of the enriched region, defined as positions where the smoothed coverage exceeded 20% of the maximum signal within the retron locus.

**Supplementary Figure S2.**
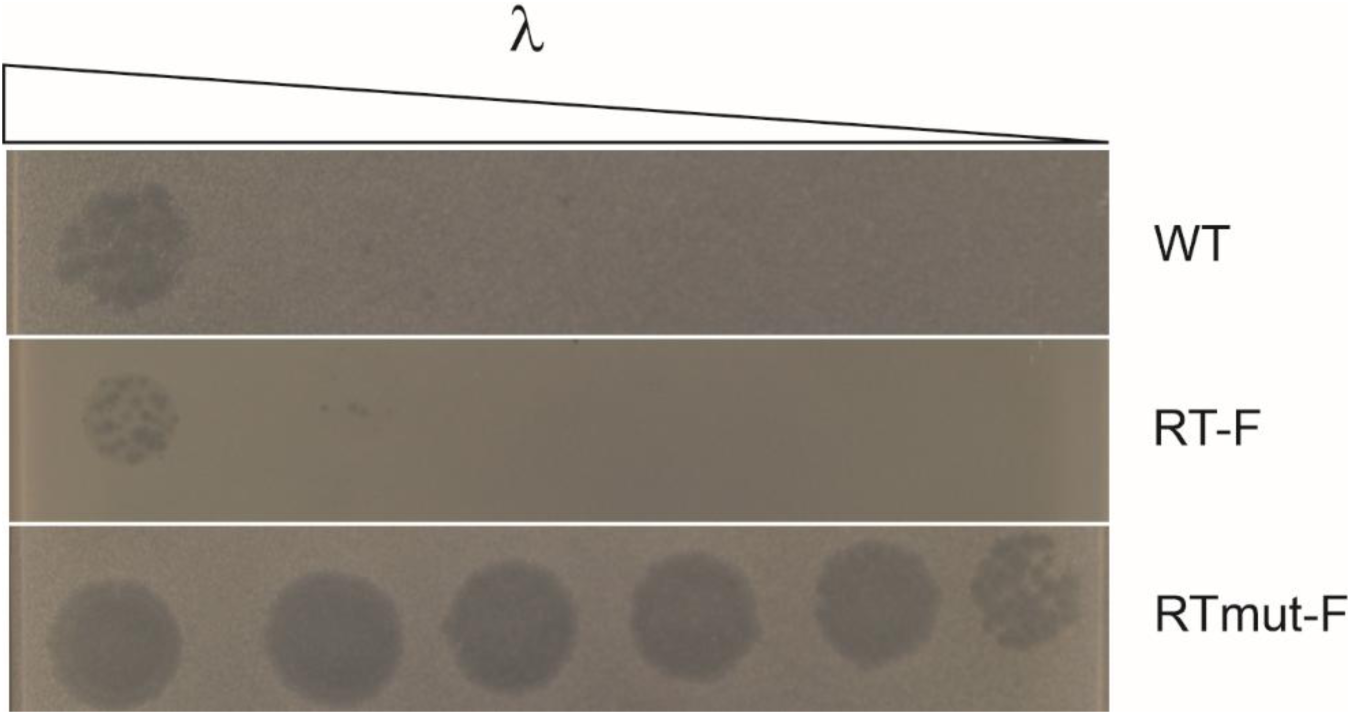
Functional validation of the C-terminally FLAG-tagged RT used in this study. Plaque assay showing the phage defense activity of Retron-Sen3 expressed in *E. coli*. Strains carried plasmids expressing the wild-type retron (WT), a variant in which the RT was fused to a C-terminal 3×FLAG tag (RT-F), or a catalytically inactive RT mutant carrying the same tag (RTmut-F; DD193–194AA). The RT-F construct retained phage defense activity comparable to the untagged retron, whereas the catalytic RT mutant failed to confer resistance, confirming that the C-terminal FLAG fusion does not impair retron function.

**Supplementary Figure S3.**
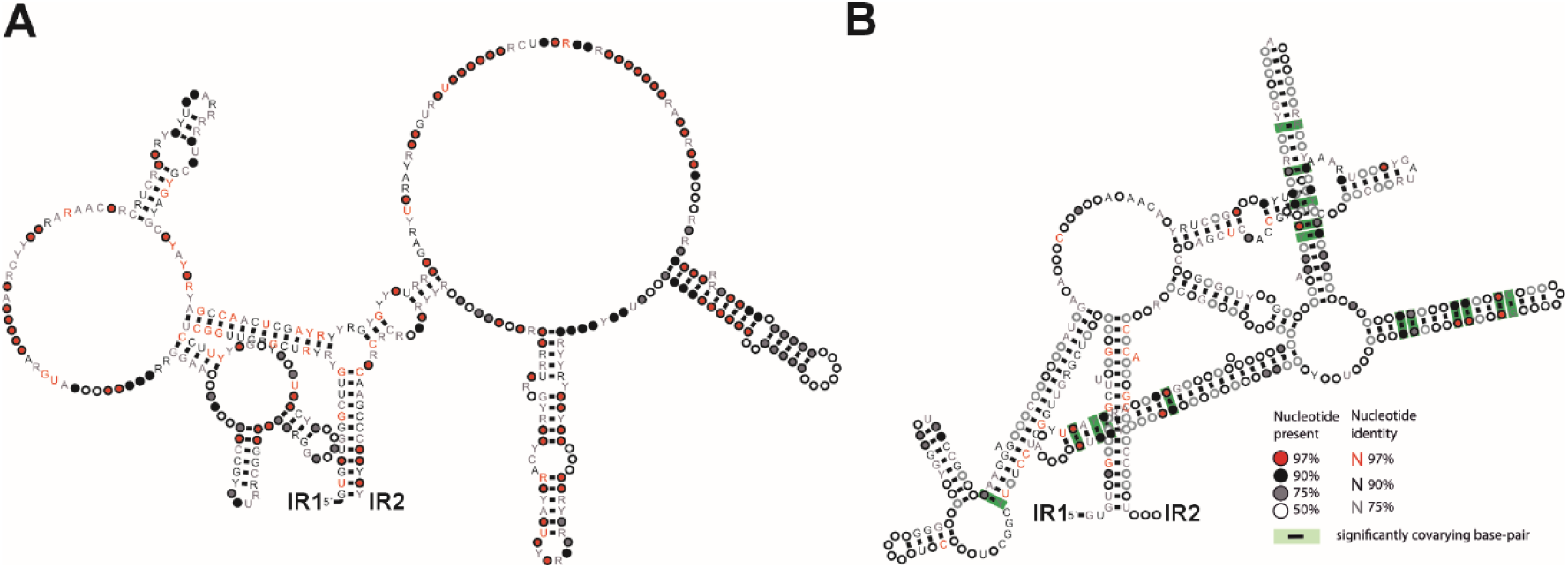
Covariance analysis of the extended transcript encompassing IR1 and IR2 in cl2075-associated retrons. **(A)** R-scape R2R representation of the consensus secondary structure of the IR1–IR2 interval derived from structure-guided alignment of 53 cl2075-associated retrons, showing nucleotide conservation at each position using IUPAC ambiguity codes. The SP ribosome-binding site (RBS) and start codon (AUG) are highlighted with a box. **(B)** CaCoFold consensus secondary structure of the IR1–IR2 interval derived from the structure-guided alignment. The SP ribosome-binding site (RBS) is highlighted. Statistically significant covarying base pairs within the SP coding region are indicated (18 covarying pairs; positive predictive value = 100%). The IR1–IR2 long-range stem is shown but does not display detectable covariation.

## Supplementary Tables

**Table S1.** Plasmid used in this study

**Table S12.** Mutations identified in phage λ escape mutants that bypass Retron-Sen3-mediated defense. All escape mutants carry single-nucleotide substitutions or frameshift insertions exclusively in the exo gene encoding the exonuclease of the λ Red recombination operon. Mutation coordinates refer to positions in the phage λ genome (GenBank accession NC_001416).

**Table S3.** Domain and subdomain organization of Sen3 SP and homologous proteins. Sequences corresponding to the N-terminal region, core domain, hydrophobic core, hydrophobic nucleus, and C-terminal amphipathic helix of Sen3 SP and its orthologs from *Yersinia* and *Xenorhabdus*, compared with the toxin Vpa2. Coordinates are indicated according to the Sen3 reference sequence. Domain boundaries were defined based on Kyte–Doolittle hydrophobicity profiles and hydrophobic moment (μH) analyses of the C-terminal region.

## Supplementary data

**Supplementary Data 1**. Newick-format phylogenetic tree of Type VI retron reverse transcriptases. The tree corresponds to the maximum-likelihood phylogeny (consensus tree) shown in Figure 3 and includes all 292 RT sequences analyzed in this study. Type IV and Type V RTs were included as outgroups for rooting. Branch support values correspond to ultrafast bootstrap replicates generated with IQ-TREE2.

