## Supplementary figures and images for "Post-translational toxin–antitoxin control and RecBCD surveillance underlie phage defense in a constitutively armed Type VI retron"

### Figure S1

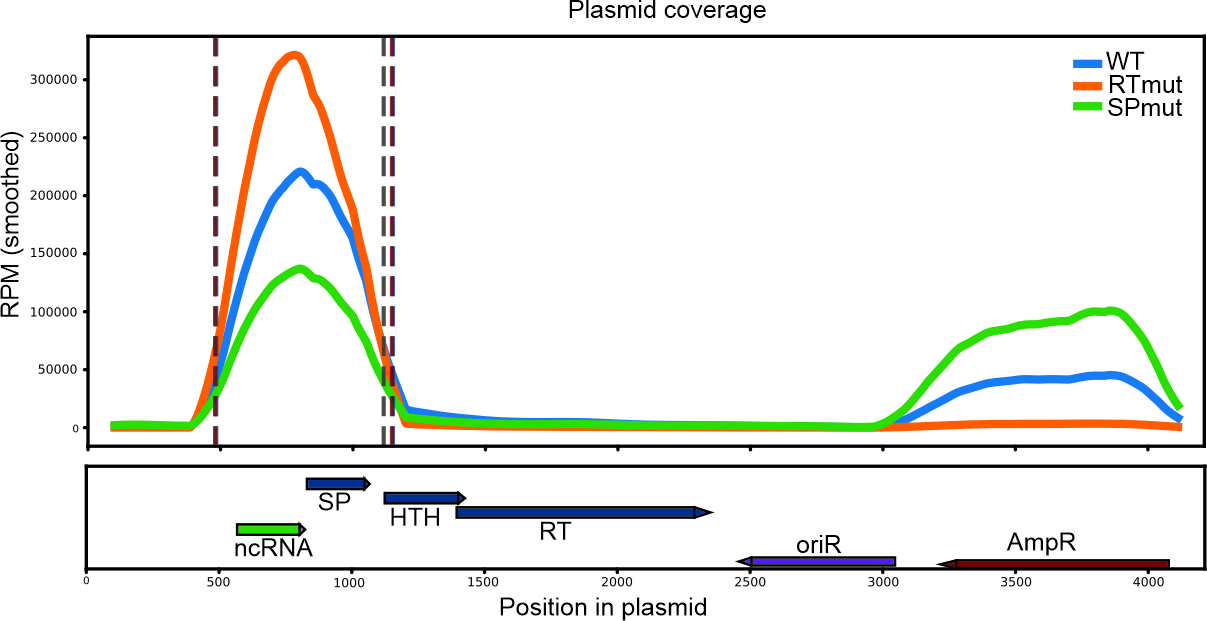

### Figure S2

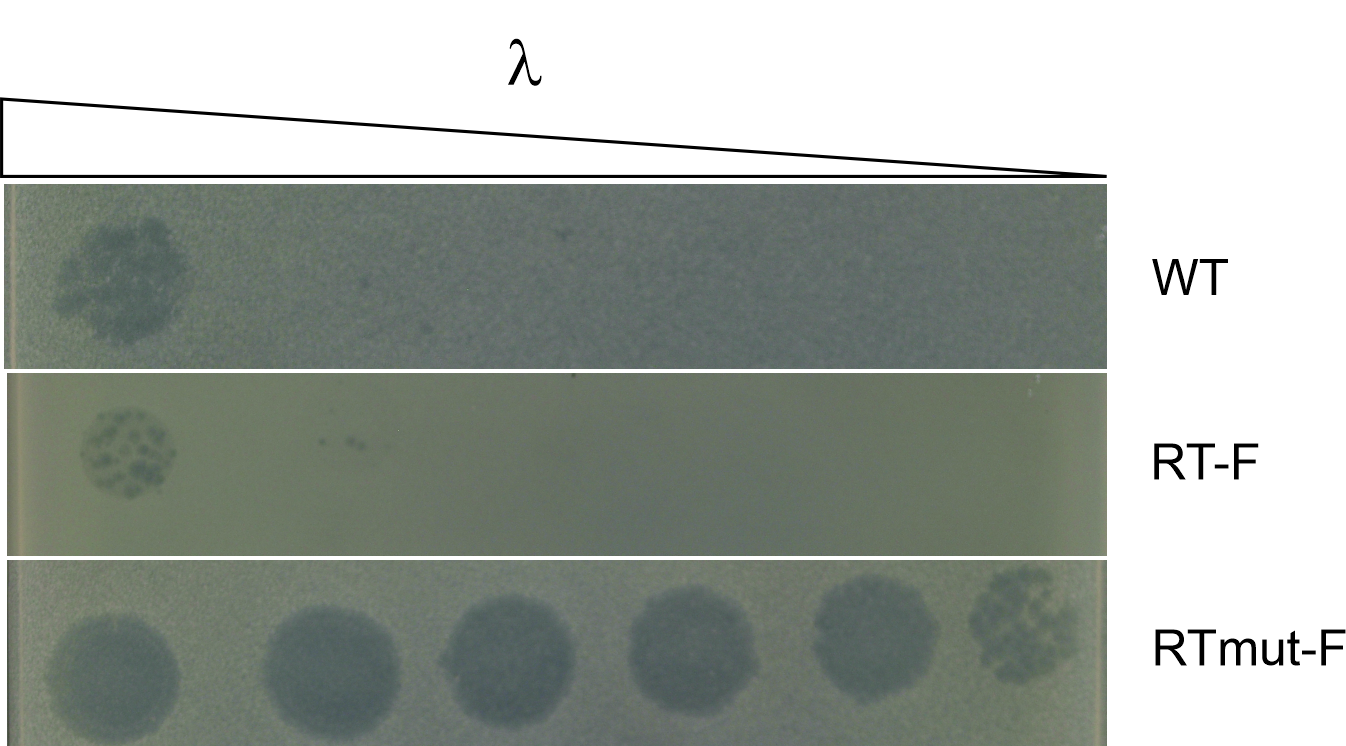

### Figure S3

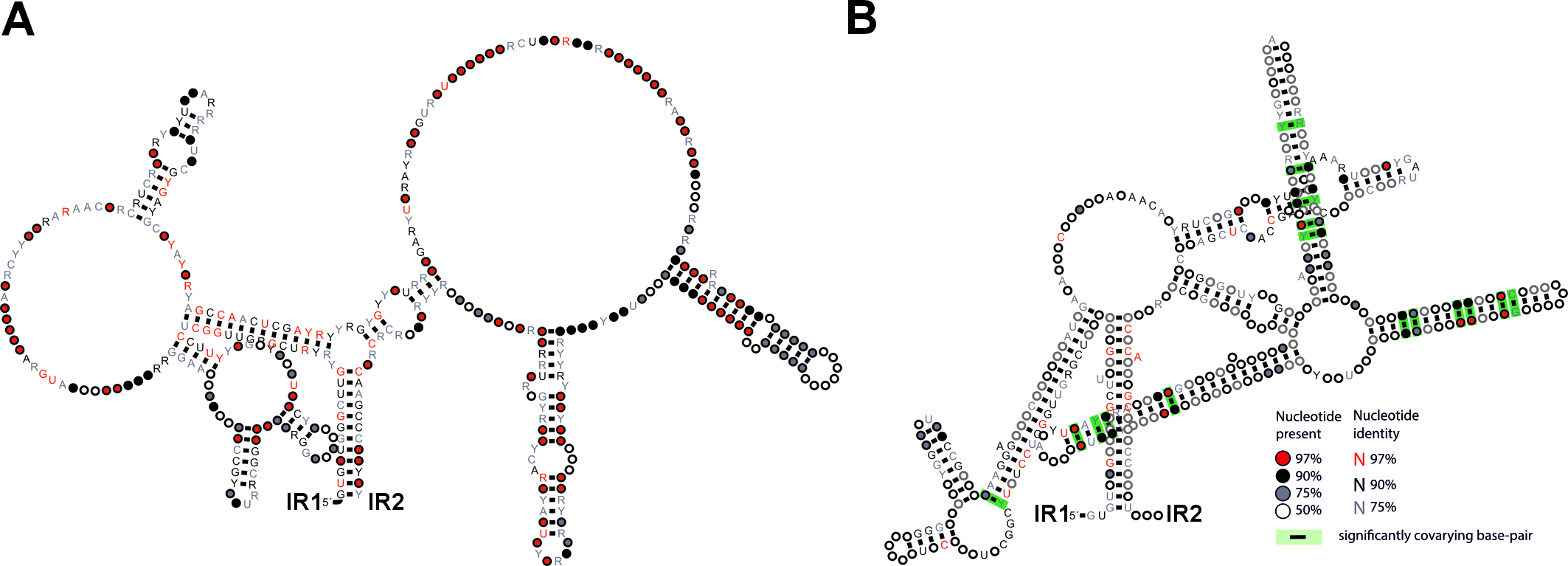
