## Supplementary material for "Post-translational toxin–antitoxin control and RecBCD surveillance underlie phage defense in a constitutively armed Type VI retron": Table S1

**Supplementary Table S1.** Plasmids used in this study

| Plasmid | Selection | Description | Source |
|---|---|---|---|
| p57m VI | Ampicillin | Retron-Sen3 locus from *S. enterica* cloned into pUC57mini | This study |
| p57m VI RTmut | Ampicillin | Derivative of p57m VI carrying the DD193–194AA substitution in the RT catalytic site | This study |
| p57m VI RTmut-F | Ampicillin | Derivative of p57m VI RTmut with C-terminal 3×FLAG tag on RT | This study |
| p57m VI RT-F | Ampicillin | Derivative of p57m VI with wild-type RT C-terminally tagged with 3×FLAG | This study |
| p57m VI SP-CF | Ampicillin | Derivative of p57m VI with SP protein C-terminally tagged with 3×FLAG | This study |
| p57m VI SPmut | Ampicillin | Derivative of p57m VI with frameshift mutation in SP (G insertion after position +36) | This study |
| p57m VI SPmut-CF | Ampicillin | Derivative of p57m VI SPmut with C-terminal 3×FLAG tag on SP | This study |
| p57m VI HTHmut | Ampicillin | Derivative of p57m VI with frameshift mutation in HTH (G insertion after position +36) | This study |
| p57m VI HTHmut-CF | Ampicillin | Derivative of p57m VI HTHmut with C-terminal 3×FLAG tag on HTH | This study |
| p57m VI Δ18nt-SL | Ampicillin | Derivative of p57m VI carrying a deletion of the 18 nucleotides comprising the 5′-conserved stem–loop of the ncRNA | This study |
| p57m VI 18nt-SLmut | Ampicillin | Derivative of p57m VI in which positions 1, 3, 5, and 8 of the 5′-conserved stem–loop are substituted. | This study |
| p57m VI ΔncRNA | Ampicillin | Derivative of p57m VI with deletion of the SL3 region of the ncRNA | This study |
| pBAD33-SP-HTH | Chloramphenicol | Wild-type SP-HTH coding sequences cloned into pBAD33 under arabinose-inducible promoter | This study |
| pBAD33-SPmut-HTH | Chloramphenicol | SP frameshift mutant with wild-type HTH cloned into pBAD33 under arabinose-inducible promoter | This study |
| pBAD33-SP-HTHmut | Chloramphenicol | Wild-type SP with HTH frameshift mutant cloned into pBAD33 under arabinose-inducible promoter | This study |
| pBAD33-gam | Chloramphenicol | Phage λ Red operon *gam* gene under arabinose-inducible promoter in pBAD33 | This study |
| pBAD33-beta | Chloramphenicol | Phage λ Red operon *beta* gene under arabinose-inducible promoter in pBAD33 | This study |
| pBAD33-exo | Chloramphenicol | Phage λ Red operon *exo* gene under arabinose-inducible promoter in pBAD33 | This study |
| pBAD33-beta+exo | Chloramphenicol | Phage λ Red operon *beta* and *exo* genes under arabinose-inducible promoter in pBAD33 | This study |
| pBAD33-gam+beta+exo | Chloramphenicol | Complete phage λ Red operon (*gam*, *beta*, *exo*) under arabinose-inducible promoter in pBAD33 | This study |
