## Supplementary material for "Post-translational toxin–antitoxin control and RecBCD surveillance underlie phage defense in a constitutively armed Type VI retron": Table S2

**Supplementary Table S2.** Mutations identified in phage λ escape mutants that bypass Retron-Sen3-mediated defense. Mutation coordinates refer to positions in the phage λ genome (GenBank accession NC_001416).

| Phage | Mutant | Mutated gene | Function | Mutation coordinate | Mutation | Effect on protein |
|---|---|---|---|---|---|---|
| λ | 1 | exo | Exonuclease | 31863 | A to G | Ser to Pro |
| | 2 | exo | Exonuclease | 31356 | A to T | Trp to Arg |
| | 3 | exo | Exonuclease | 31972 | +C | Frameshift |
| | 4 | exo | Exonuclease | 34531 | +T | Frameshift |
| | 5 | exo | Exonuclease | 31676 | G to A | Pro to Leu |
