## Supplementary material for "Post-translational toxin–antitoxin control and RecBCD surveillance underlie phage defense in a constitutively armed Type VI retron": Table S3

**Supplementary Table S3. Domain and subdomain organization of Sen3 SP and homologous proteins.**

| Feature | Sen3 coordinates (aa) | Sen3 sequence | Yersinia sequence | Xenorhabdus sequence | Vpa2 sequence | Notes |
|---|---|---|---|---|---|---|
| **N-terminal region** | 1–25 | MKYETLRNTVATLQKLRDVHYSQLD | MNNQTLKNSIATLQSLRDAHYSQLD | MKHDTLKNTIATLEKLRDAHCSQLD | NNTQ | *Polar extension absent in Vpa2* |
| **Core domain** | 26–60 | AGALAELDDVLQQLRNMLDGSERRELEHGE | DASALAELDEVLLQLRKLPKGPEVHKSSHGELV | DAGALAELDDVLQQLRSHQDSPVKSGNRSDDI | MSNNTQRDQQVNIIKPTKREKRQKQFKRFLKALSW | *Conserved structural scaffold* |
| **Hydrophobic core** | 47–69 | ELVFRALRITDIAIRAVSNLTDW | ELVLRVFRIIDIILRLVSNIN | VFRALRIIDIVLRLVTNITD | SWLYRIWYWFNFLLGLFD | *Apolar region preceding the C-terminal helix* |
| **Hydrophobic nucleus** | 63–69 | RITDIAI | RIIDIIL | RIIDIVL | IWYWFNF | *Peak KD; aromatic enrichment in Vpa2* |
| **Amphipathic helix** | 63–75 | RITDIAIRAVSNLT | RIIDIILRLVSNIN | RIIDIVLRLVTNIT | – | *Amphipathic in Sen3, Yersinia and Xenorhabdus; absent in Vpa2* |
| **C-terminal helix** | 61–79 | LRITDIAIRAVSNLTDWMK | FRIIDIILRLVSNINDWMK | LRIIDIVLRLVTNITDWMK | RIWYWFNFLLGLFDGGDT | *Conserved helix; non-amphipathic in Vpa2* |
